# Single-cell RNA-seq reveals spatially restricted multicellular fibrotic niches during lung fibrosis

**DOI:** 10.1101/569855

**Authors:** Nikita Joshi, Satoshi Watanabe, Rohan Verma, Renea P. Jablonski, Ching-I Chen, Paul Cheresh, Paul A. Reyfman, Alexandra C. McQuattie-Pimentel, Lango Sichizya, Annette S. Flozak, Cara J. Gottardi, Carla M. Cuda, Harris Perlman, Manu Jain, David W. Kamp, GR Scott Budinger, Alexander V. Misharin

## Abstract

Ontologically distinct populations of macrophages differentially contribute to organ fibrosis through unknown mechanisms. We applied lineage tracing, spatial methods and single-cell RNA-seq to a spatially-restricted model of asbestos-induced pulmonary fibrosis. We demonstrate that while tissue-resident interstitial macrophages, tissue-resident alveolar macrophages, and monocyte-derived alveolar macrophages are present in the fibrotic niche, only monocyte-derived alveolar macrophages are causally related to fibrosis. Monocyte-derived alveolar macrophages were specifically localized to fibrotic regions in the proximity of fibroblasts where they expressed molecules known to drive fibroblast proliferation, including PDGFA. Moreover, we identified autocrine M-CSF/M-CSFR signaling in monocyte-derived alveolar macrophages as a novel mechanism promoting their self-maintenance and persistence in the fibrotic niche. Pharmacological blockade of M-CSF signaling led to disappearance of the established population of monocyte-derived alveolar macrophages. Thus, our data indicate that monocyte-derived alveolar macrophages are specifically recruited to the fibrotic niche where they are maintained by autocrine signaling and drive fibrosis by stimulating fibroblast proliferation.

## Introduction

Pulmonary fibrosis is a complex process that is clinically characterized by a progressive increase in the number and size of spatially restricted areas of fibrosis (Raghu et al., 2018). Indeed, the three dimensional distribution of these lesions on chest computed tomography combined with radiographic features of the fibrotic regions are critical to the diagnosis and classification of pulmonary fibrosis. Single-cell RNA-seq offers the opportunity to examine interactions between cell populations within these areas of fibrosis, but the process of tissue dissociation precludes understanding the spatial relationships between the cells (Reyfman et al., 2018). We reasoned that a combination of genetic lineage tracing, single-cell RNA-seq, and spatial methods could be combined with genetic or pharmacologic interventions to identify conserved intracellular signaling events that might promote or sustain multicellular fibrotic niches in the lung. To test this hypothesis, we used a model of asbestos-induced lung fibrosis. Exposure to asbestos can induce the development of pulmonary fibrosis years after the exposure has ceased (Cugell and Kamp, 2004), and historic and ongoing exposure to asbestos fibers remains an important occupational cause of pulmonary fibrosis. After inhalation, asbestos fibers remain lodged in small airways in rodents (Adamson and Bowden, 1987a; Adamson and Bowden, 1987b; Roggli et al., 1987) creating spatially restricted regions of lung fibrosis.

Recently, investigators have identified several distinct macrophage populations in the lung. Lung tissue-resident interstitial macrophages have been shown to include perivascular and peribronchial macrophages, which demonstrate distinct anatomic localization and function (Gibbings et al., 2017; Lim et al., 2018; Loyher et al., 2018). Tissue-resident alveolar macrophages originate from fetal monocytes, populate the alveolar niche soon after birth, are capable of self-renewal, and, in mouse models, persist in the lung without appreciable input from myeloid cells for up to 14 months (Guilliams et al., 2013; Janssen et al., 2011; Misharin et al., 2017; Yona et al., 2013). In response to alveolar macrophage depletion and/or injury, monocytes are recruited to the lung where factors present in the microenvironment drive their differentiation into alveolar macrophages (Gibbings et al., 2015; Lavin et al., 2014). Alveolar macrophages were recognized to play a critical role in asbestos-induced injury when Dostert et al. reported that potassium currents in alveolar macrophages attempting to engulf asbestos fibers resulted in activation of the NLRP3 inflammasome (Dostert et al., 2008), which is essential for the development of asbestos-mediated lung fibrosis (dos Santos et al., 2015). Whether one or more of these macrophage populations is necessary for the development of fibrosis in response to asbestos remains unclear.

Here we used lineage tracing and a genetic deletion strategy to show that while tissue-resident interstitial macrophages, tissue-resident alveolar macrophages, and monocyte-derived alveolar macrophages are present in the fibrotic niche, only monocyte-derived alveolar macrophages are causally related to fibrosis. Spatial methods demonstrate that monocyte-derived alveolar macrophages, expressing profibrotic genes causally linked to fibrosis, form pathogenic multicellular niches with injured epithelial cells and tissue fibroblasts. Moreover, single-cell transcriptomic analysis of lungs from mice treated with asbestos or bleomycin suggested that monocyte-derived alveolar macrophages are sustained within the fibrotic niche by autocrine signaling through M-CSF/M-CSFR, which was confirmed by administration of blocking antibodies. These changes were largely confirmed in single-cell transcriptomic data from patients with pulmonary fibrosis. Our findings suggest how single-cell RNA-seq data from multiple laboratories can be combined and validated to identify conserved intercellular interactions within multicellular fibrotic microdomains that might be targeted for the treatment of pulmonary fibrosis.

## Results

### Recruitment of monocyte-derived alveolar macrophages distinguishes the response to asbestos from a non-fibrogenic particle

We quantified monocyte and macrophage populations in the mouse lung via flow cytometry 14 days after the intratracheal administration of asbestos, which reproducibly induces lung fibrosis, or a standard control particle TiO_2_ that does not induce fibrosis (both 100 μg) (see Figure S1A for gating strategy). We found that the number of monocytes and alveolar macrophages (CD64^+^Siglec F^+^) were similarly increased in mice administered either asbestos or TiO_2_ when compared with naïve mice, while non-alveolar macrophages (CD64^+^Siglec F^−^) were increased only in asbestos-treated mice (Figure 1A). While the increase in alveolar macrophages was largely attributable to expansion of a Siglec F^high^ population in the TiO_2_-treated animals, the increase in alveolar macrophages in animals administered asbestos was attributable to both an expansion in Siglec F^high^ alveolar macrophages and the emergence of a population of Siglec F^low^ alveolar macrophages (∼15% of the total alveolar macrophage pool) (Figure 1A-C).

**Figure 1.**
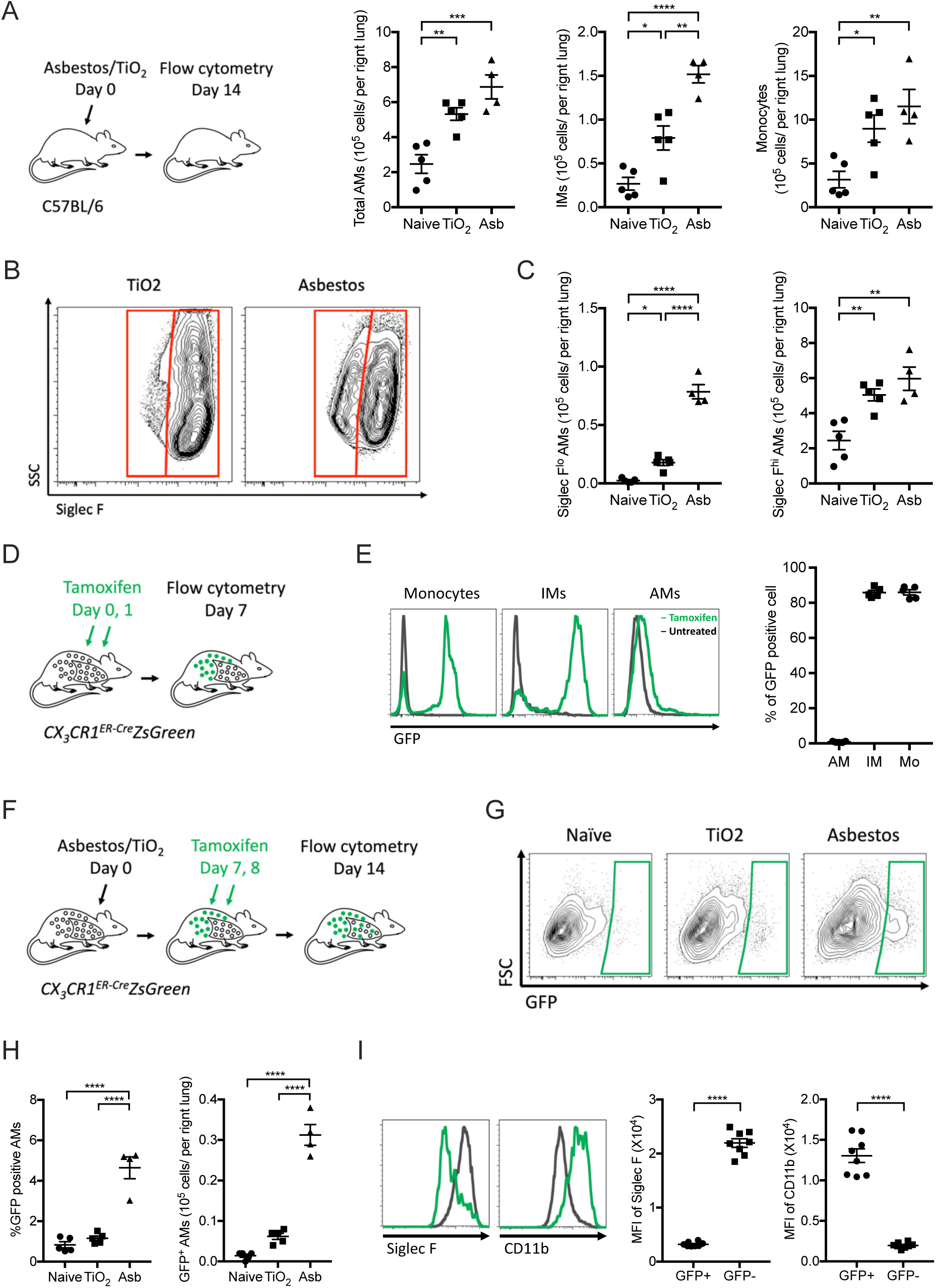
Exposure to asbestos or TiO_2_ is distinguished by the recruitment of monocyte-derived alveolar macrophages to the lung. **A**. Mice were administered crocidolite asbestos or TiO_2_ (both at 100 μg, intratracheally) and monocyte and macrophage populations were quantified by flow cytometry 14 days later (See Figure S1A,B for gating strategy and quantification of other myeloid cell populations). **B**. Representative contour plots gated on alveolar macrophages (CD64^+^Siglec F^+^) from asbestos-treated or TiO_2_-treated animals. **C**. Quantification of Siglec F^low^ and Siglec F^high^ alveolar macrophages from naïve, TiO_2_-or asbestos-exposed animals according to gating in panel B. **D**. Schematic of the experimental design in E. **E**. *Cx3cr1*^*ER-Cre*^*ZsGreen* mice were treated with tamoxifen and percentage of GFP-positive classical monocytes, interstitial macrophages and alveolar macrophages was assessed by flow cytometry (representative histograms showing GFP expression are shown, black line – control, green line – tamoxifen-treated). **F.** Schematic of experimental design for G-I: lineage tracing system to track the ontogeny of alveolar macrophages after intratracheal administration of asbestos or TiO_2_. *Cx3cr1*^*ER-Cre*^*ZsGreen* mice were treated with asbestos or TiO_2_ and tamoxifen was administered as two boluses at days 7 and 8. The number of GFP+ alveolar macrophages was analyzed 7 days later. **G-H**. Representative contour plots (**G**) and quantification (**H**) of GFP-positive monocyte-derived alveolar macrophages after asbestos or TiO_2_ exposure. **I**. Representative histograms and median fluorescence intensity (MFI) demonstrating expression of Siglec F and CD11b on monocyte-derived alveolar macrophages 14 days after exposure to asbestos. All data presented as mean±SEM, 4–5 mice per group, one-way ANOVA with Tukey-Kramer test for multiple comparisons; *, P < 0.05; **, P < 0.01; ***, P < 0.001; ****, P < 0.0001. Representative data from two independent experiments is shown.

**Figure S1.**
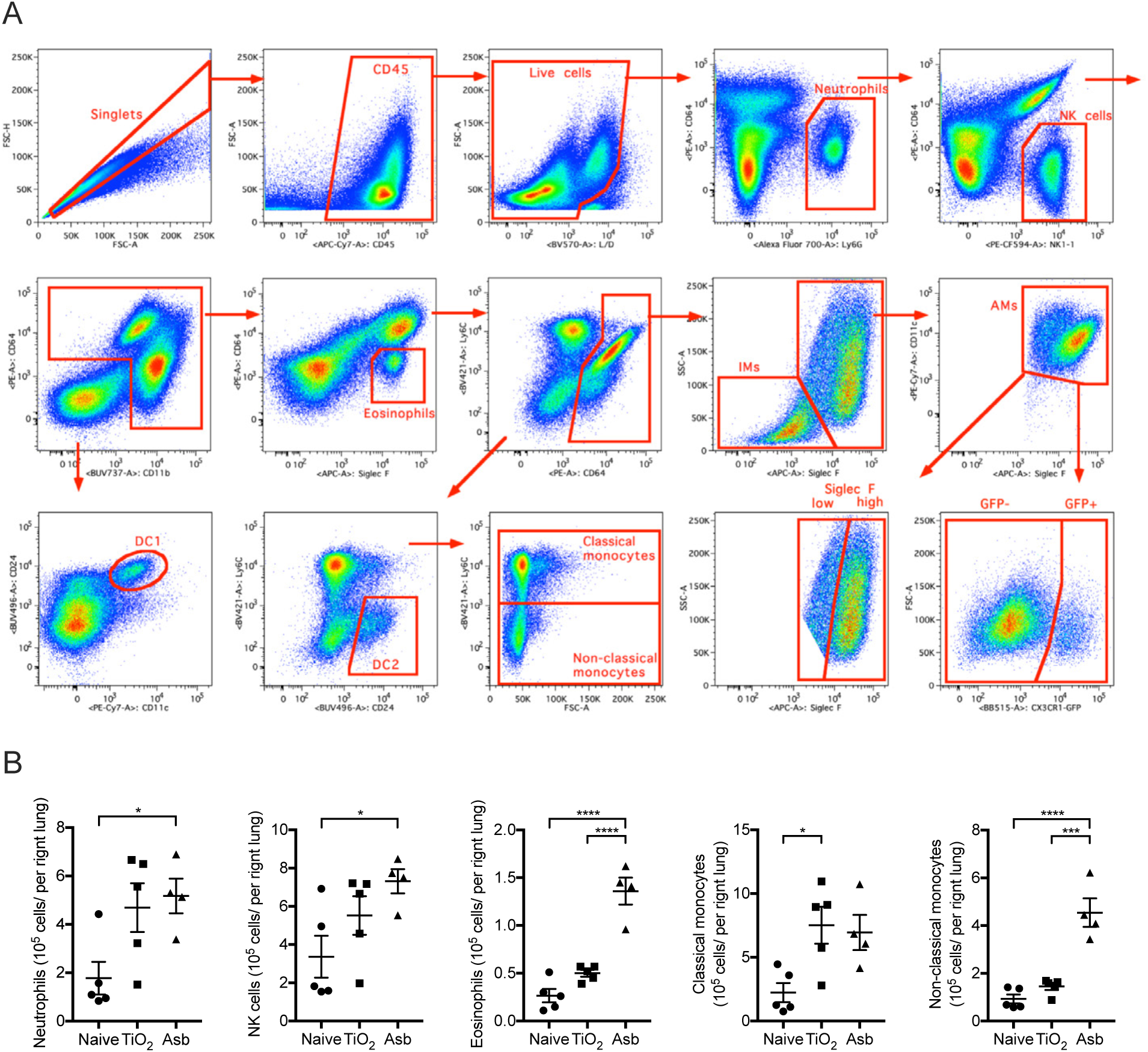
(refers to Figure 1). **A.** Gating strategy for quantification of myeloid populations in the lung. **B.** Quantification of myeloid cell populations by flow cytometry 14 days after mice were administered crocidolite asbestos or TiO_2_ (both at 100 μg, intratracheally). All data presented as mean±SEM, 4–5 mice per group, one-way ANOVA with Tukey-Kramer test for multiple comparisons; *, P < 0.05; **, P < 0.01; ****, P < 0.0001. Representative data from two independent experiments is shown.

We used a genetic lineage tracing system to determine whether Siglec F^low^ alveolar macrophages that were expanded in asbestos-treated animals but absent in TiO_2_-treated animals were derived from the recruitment of monocytes or the expansion of tissue-resident alveolar macrophages. Circulating Ly6C^high^ classical monocytes express high levels of *Cx3cr1*, while tissue-resident alveolar macrophages do not (Gibbings et al., 2017; Yona et al., 2013). Accordingly, we crossed *Cx3cr1*^*ER-Cre*^ animals to ZsGreen reporter mice to track the fate of monocyte-derived cells in the lung during asbestos-mediated fibrosis (Figure 1D). In this system, 7 days after tamoxifen administration 86.8% of circulating monocytes were GFP-positive (Figure 1E), but because they are short-lived, they disappeared after tamoxifen pulse (Yona et al., 2013). However, if GFP-labeled monocytes give rise to long-living macrophages, such as monocyte-derived alveolar macrophages, they will be permanently labeled with GFP, even after the expression of endogenous *Cx3cr1* has ceased. As expected, long-lived tissue-resident alveolar macrophages, which do not express *Cx3cr1*, were not labeled in this system (Figure 1E). In contrast, 86.7% of the tissue-resident peribronchial and perivascular interstitial macrophages, which also express *Cx3cr1*, become GFP-positive after tamoxifen pulse (Figure 1E).

To assess the contribution of monocyte-derived macrophages to the expansion of the alveolar macrophage pool, the reporter mice were treated with tamoxifen via oral gavage 7 and 8 days after the administration of asbestos or TiO_2_ and lungs were analyzed by flow cytometry at day 14 (Figure 1F). Administration of asbestos resulted in an influx of GFP^+^ alveolar macrophages, which were Siglec F^low^ and CD11b^high^ (Figure 1G-I). In total, 4.9% of monocyte-derived alveolar macrophages (CD64^+^Siglec F^low^) were GFP positive one week after tamoxifen pulse in asbestos-exposed mice (Figure 1H-I). In contrast, in mice treated with TiO_2_ only 1.4% of alveolar macrophages were GFP-positive, which was comparable to naïve mice (Figure 1H-I).

### Monocyte-derived alveolar macrophages and not tissue-resident interstitial macrophages contribute to the macrophage pool in the fibrotic niche

We took advantage of the GFP label on monocyte-derived alveolar macrophages in our lineage tracing system to determine whether recruitment of these cells in response to asbestos was spatially restricted to areas near the fibers. Fibrosis was restricted to regions of bronchoalveolar duct junctions where asbestos fibers had lodged, but was absent in the distal lung (Figure 2A). Immunofluorescent microscopy demonstrated that GFP^+^ cells were specifically found in these fibrotic areas and both GFP^+^ and GFP^−^ cells appeared to engulf asbestos fibers (Figure 2B, C). These cells are located in the alveolar space and express MerTK and Siglec F confirming their identity as monocyte-derived alveolar macrophages (Figure S2).

**Figure 2.**
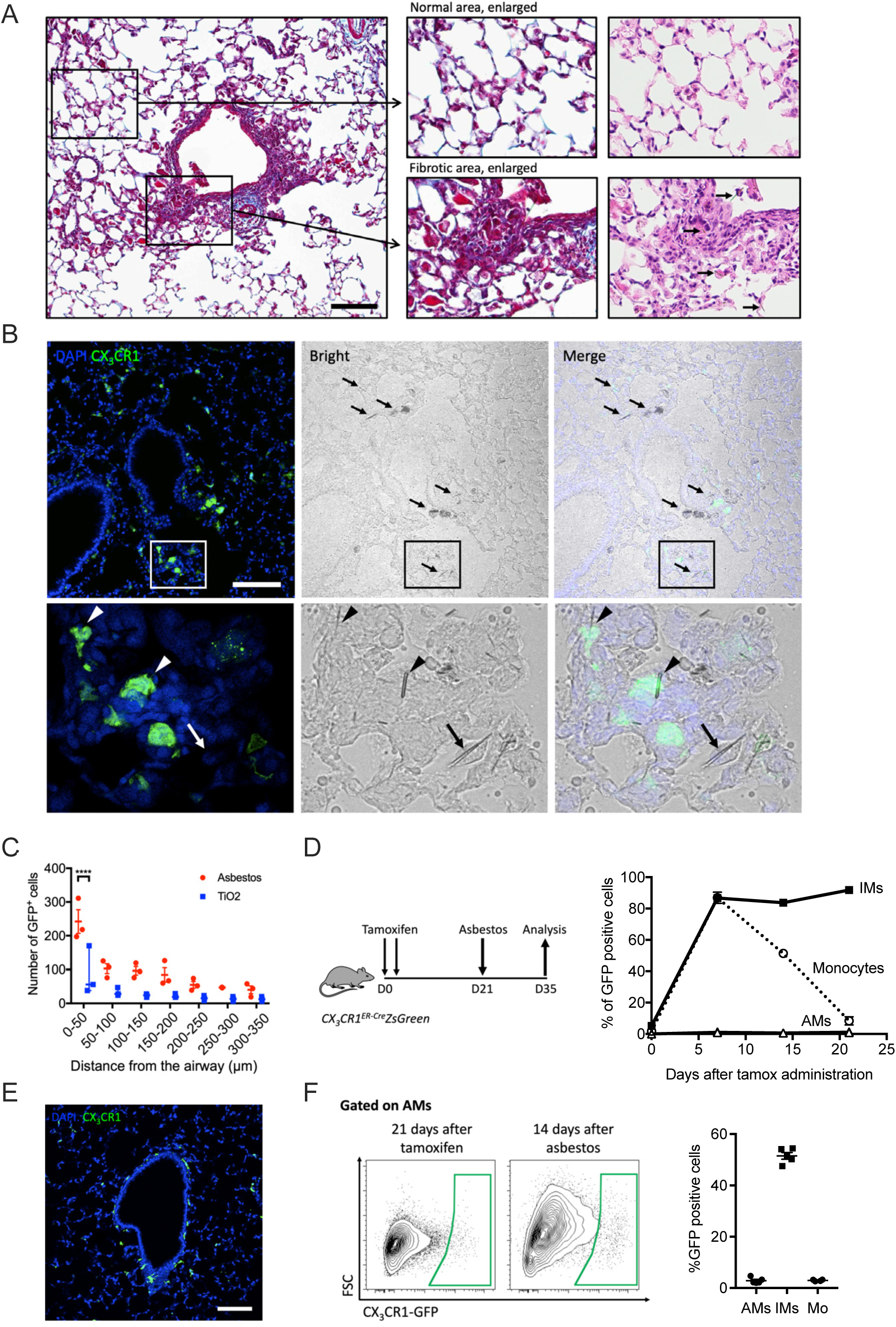
Recruitment of monocyte-derived alveolar macrophages is spatially restricted to areas near asbestos fibers. **A.** The intratracheal administration of asbestos fibers induces fibrosis near bronchoalveolar duct junctions where asbestos fibers lodge. Left panel shows low power image of a medium-sized airway (Mason’s trichrome, scale bar 100 µm), right panel shows high power images (Mason’s trichrome and H&E, respectively). Areas of fibrosis develop adjacent to the airway in which asbestos fibers can be observed (arrows, bottom right panel). In contrast alveolar structures in the distal lung parenchyma are relatively preserved. **B.** Top panel: representative lung histology from *Cx3cr1*^*ER-Cre*^*ZsGreen* mice treated with tamoxifen on day 14 and 15 (10 mg, via oral gavage) and harvested 21 days after asbestos exposure (scale bar 100 µm). Bottom panels correspond to areas outlined in boxes. Left panels: monocyte-derived cells are GFP-positive, nuclei stained with DAPI. Middle panels: phase contrast images, asbestos fibers are indicated by arrows. Right panels: merge. Bottom panels: asbestos fibers are surrounded by GFP-positive cells (arrowheads) and GFP negative cells (arrow) with macrophage morphology. **C**. Quantification of GFP-positive monocyte-derived alveolar macrophages in peribronchial regions in asbestos-and TiO_2_-treated mice (two-way ANOVA with Tukey’s multiple comparisons test, p=0.0032). **D**. Schematic of experimental design, and kinetics of GFP-positive monocytes, tissue-resident interstitial macrophages and tissue-resident alveolar macrophages after tamoxifen pulse in naïve *Cx3cr1*^*ER-Cre*^*ZsGreen* mice. Percentage of GFP-positive cells was assessed by flow cytometry. **E.** Representative fluorescent images showing GFP-positive tissue-resident interstitial macrophages 21 days after tamoxifen pulse in naïve animals. **F.** Representative contour plots showing GFP expression in alveolar macrophages from *Cx3cr1*^*ER-Cre*^*ZsGreen* mice 21 days after tamoxifen and 14 days after asbestos instillation. Percentage of GFP-positive classical monocytes, interstitial macrophages and alveolar macrophages was assessed by flow cytometry according to gate in F. All data presented as mean±SEM, 3–5 mice per group or time-point.

**Figure S2.**
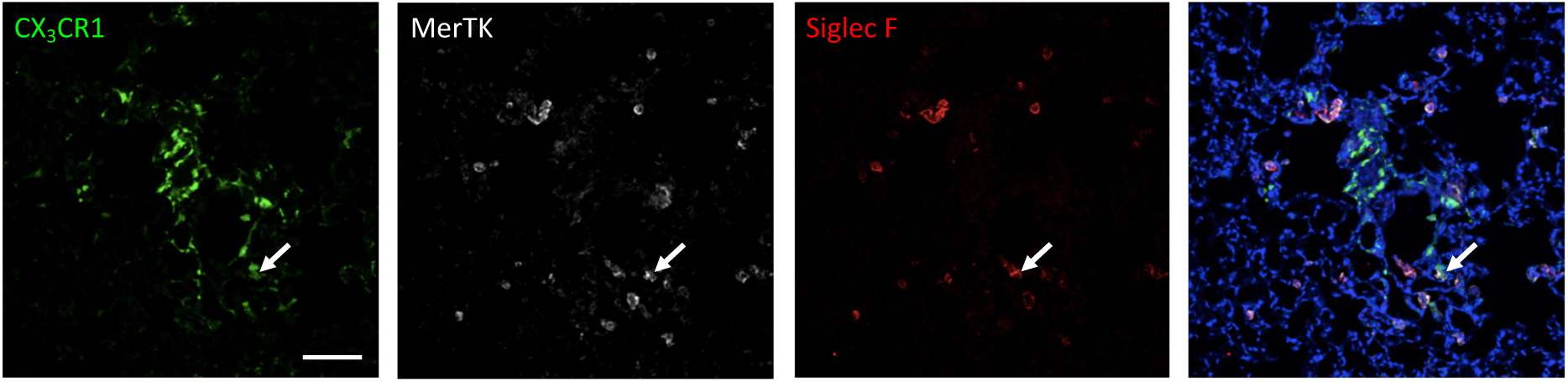
Monocyte-derived alveolar macrophages express canonical alveolar macrophage markers after asbestos exposure. (refers to Figure 2). Representative fluorescent images showing expression (arrows) of CX_3_CR1-GFP (green), MerTK (white), and Siglec F (red). Note, CX_3_CR1-GFP^+^Siglec F^−^ macrophages surrounding the airway in the center of the image. Scale bar 100 μm.

Tissue-resident peribronchial interstitial macrophages (CD64^+^Siglec F^−^) have recently been reported to contribute to the population of tumor-associated macrophages (Loyher et al., 2018). To exclude the possibility that these tissue-resident interstitial macrophages can migrate to the alveolar space in response to asbestos-induced epithelial injury and differentiate into alveolar macrophages, we treated reporter mice with tamoxifen to label tissue-resident interstitial macrophages (Figure 2D). By 21 days after tamoxifen treatment, virtually all GFP-labeled monocytes disappeared from circulation while 82% of tissue-resident interstitial macrophages remained GFP-positive (Figure 2D-E). These mice were treated with asbestos intratracheally and the lungs were analyzed by flow cytometry 14 days later (Figure 2D). The percentage of GFP-positive alveolar macrophages in asbestos-exposed mice was not different from non-exposed mice, suggesting that monocytes, and not tissue-resident interstitial macrophages drive the expansion of alveolar macrophages at the sites of asbestos-mediated injury and fibrosis (Figure 2F).

### Genetic deletion of monocyte-derived alveolar macrophages attenuates asbestos-induced pulmonary fibrosis

Previously, we have shown that the loss of caspase-8 in monocyte-derived alveolar macrophages results in their death via necroptosis during differentiation (Misharin et al., 2017). Accordingly, we used macrophage-specific deletion of *Casp8* with or without prevention of necroptosis through deletion of *Ripk3* to determine whether monocyte-derived alveolar macrophages are necessary for the development of asbestos-induced pulmonary fibrosis. Control mice (*Casp8*^*flox/flox*^) and mice lacking caspase-8 in alveolar macrophages (*CD11c*^*Cre*^*Casp8*^*flox/flox*^) were administered either asbestos or TiO_2_ intratracheally. The number of monocyte-derived alveolar macrophages (CD64^+^Siglec F^low^) was significantly reduced in *CD11c*^*Cre*^*Casp8*^*flox/flox*^ compared with *Casp8*^*flox/flox*^ controls 28 days after the administration of asbestos (Figure 3A-D). Other cell populations were unchanged in the *CD11c*^*Cre*^*Casp8*^*flox/flox*^ mice compared to *Casp8*^*flox/flox*^ controls with the exception of eosinophils, which were reduced in the *CD11c*^*Cre*^*Casp8*^*flox/flox*^ mice compared with the *Casp8*^*flox/flox*^ controls (Figure 3E; Figure S3A). *CD11c*^*Cre*^*Casp8*^*flox/flox*^ mice showed less asbestos-induced fibrosis compared with the *Casp8*^*flox/flox*^ controls (Figure 3F-H). To confirm that the protection from fibrosis in *CD11c*^*Cre*^*Casp8*^*flox/flox*^ mice resulted from necroptotic loss of monocyte-derived alveolar macrophages, we performed the same experiments in *CD11c*^*Cre*^*Casp8*^*flox/flox*^*Ripk3*^*-/-*^ mice and *Ripk3*^*-/-*^ mice. While the global loss of *Ripk3* did not affect the recruitment of monocyte-derived alveolar macrophages or the severity of fibrosis, the loss of *Ripk3* in *CD11c*^*Cre*^*Casp8*^*flox/flox*^*Ripk3*^*-/-*^ mice restored monocyte-derived alveolar macrophages and rescued asbestos-induced fibrosis (Figure 3A-H). To determine whether tissue-resident alveolar macrophages also contributed to the development of asbestos-induced fibrosis, we depleted tissue-resident alveolar macrophages by treating mice with intratracheal liposomal clodronate 24 hours prior to the administration of intratracheal asbestos or TiO_2_. There was no difference in fibrosis 28 days later as determined by quantitative scoring of lung sections and measurements of lung soluble collagen using picrosirius red precipitation, suggesting that tissue-resident alveolar macrophages are not required for the development of fibrosis (Figure S3B). These data demonstrate that monocyte-derived alveolar macrophages are necessary to develop a spatially-restricted fibrotic niche in response to asbestos.

**Figure 3.**
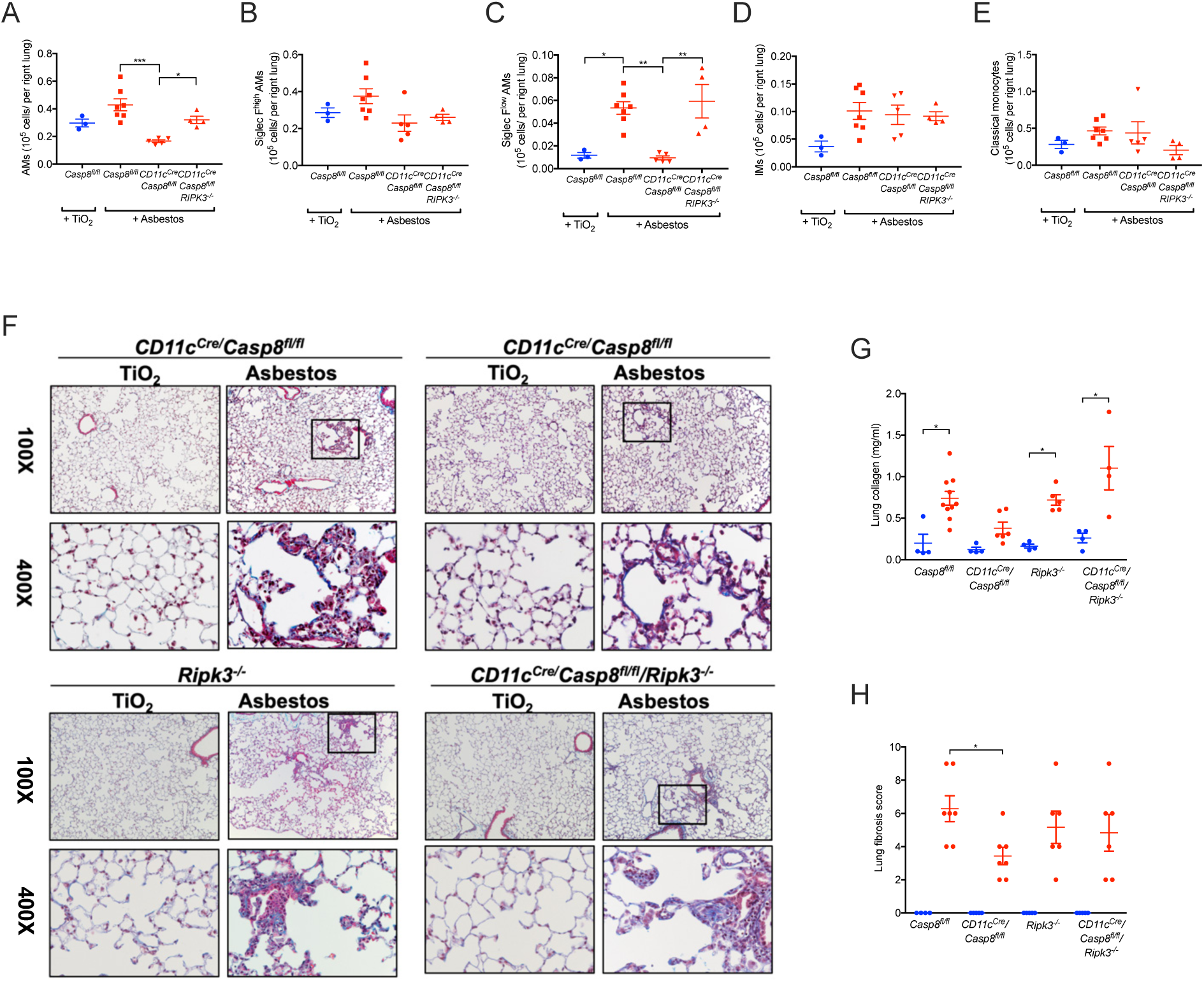
Deletion of monocyte-derived alveolar macrophages attenuates asbestos-induced pulmonary fibrosis. *Casp8*^*flox/flox*^, *CD11c*^*Cre*^*Casp8*^*flox/flox*^, *CD11c*^*Cre*^*Casp8*^*flox/flox*^*Ripk3*^*-/-*^and *Ripk3*^*-/-*^ mice were administered crocidolite asbestos or TiO_2_ (both at 100 μg, intratracheally) and the lungs were harvested 28 days later. **A-E**. Lung were analyzed using flow cytometry to quantify monocyte and macrophage populations. **F**. Representative histologic images (Mason’s trichrome, top, 100X, bottom, 400X). **G.** Quantification of soluble collagen in lung homogenates. **H.** Blinded scoring (Ashcroft score) of a single longitudinal section from each mouse. Blue colored circles refer to TiO_2_ treatment, red colored circles, squares and triangles refer to asbestos administration. All data presented as mean±SEM, 3–7 mice per group, one-way ANOVA with Tukey-Kramer test for multiple comparisons. *, p< 0.05; **, p< 0.01;***, P < 0.001.

**Figure S3.**
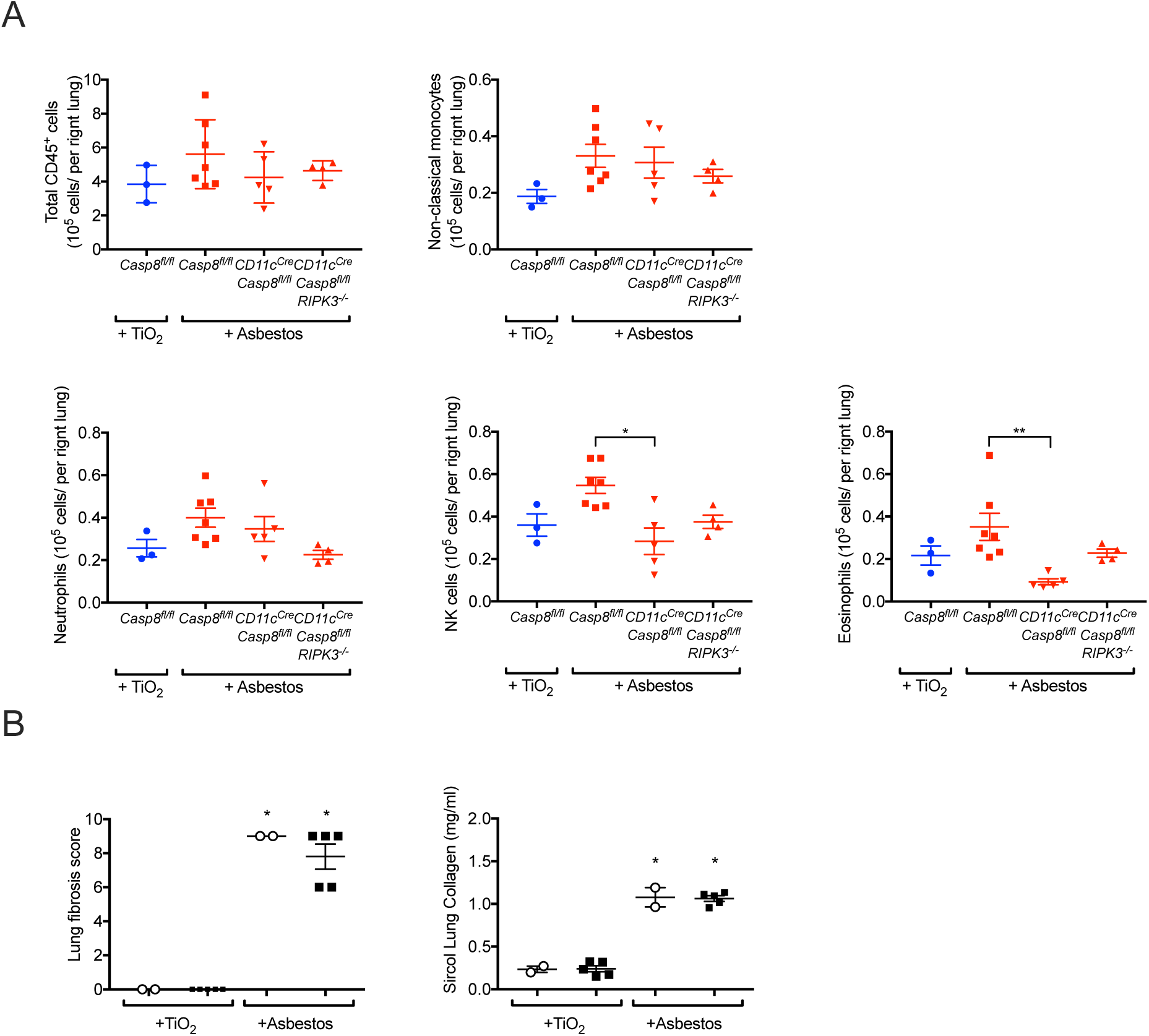
Monocyte-derived and not tissue-resident alveolar macrophages are required for the development of asbestos-induced pulmonary fibrosis (Refers to Figure 3). **A.** Quantification of myeloid cell populations by flow cytometry 28 days after mice were administered crocidolite asbestos or TiO_2_ (both at 100 μg, intratracheally). All data presented as mean±SEM, 3–7 mice per group, one-way ANOVA, with Bonferroni correction for multiple comparisons. * p<0.05. **B.** Depletion of tissue-resident alveolar macrophages does not alter the severity of asbestos-induced fibrosis. Blinded scoring of a single longitudinal section from each mouse and quantification of soluble collagen in lung homogenates. Circles refer to controls (PBS) and squares refer to treatment with clodronate-loaded liposomes. All data presented as mean±SEM, 2–5 mice per group, one-way ANOVA, with Tukey-Kramer test for multiple comparisons. *, p<0.05; **, p< 0.01.

### Single-cell RNA-seq identifies a heterogeneous response to asbestos exposure among different resident lung populations

We reasoned that single-cell RNA-seq combined with spatial techniques, such as *in situ* RNA hybridization could be used to gain insights into cellular interactions by which monocyte-derived alveolar macrophages are maintained and signal within fibrotic niches in the asbestos-exposed lung. Accordingly, we performed single-cell RNA-seq on unenriched single-cell suspensions from the lungs of wild-type mice 14 days after exposure to either asbestos or TiO_2_, when fibrosis is just beginning. After excluding doublets and low quality cells, 24,060 cells from two libraries were used for analysis and clustering was performed using Seurat R toolkit. Clusters were annotated according to cell type, based on the expression of cell type-specific genes (Figure 4A, Supplemental Table 1). We captured and resolved all major cell populations in the mouse lung, including interstitial and alveolar macrophages, alveolar epithelial type I and type II cells, dendritic cells, fibroblasts, mesothelial cells, smooth muscle cells and several subsets of endothelial cells. Each cluster included cells from both experimental groups (i.e. mice exposed to asbestos or TiO_2_) (Figure S4A-B). In agreement with our lineage-tracing studies indicating influx of monocyte-derived alveolar macrophages, we identified a subcluster of macrophages (Figure 4B) that was disproportionately represented by the cells from the asbestos-exposed animal (Figure S4B).

**Figure 4.**
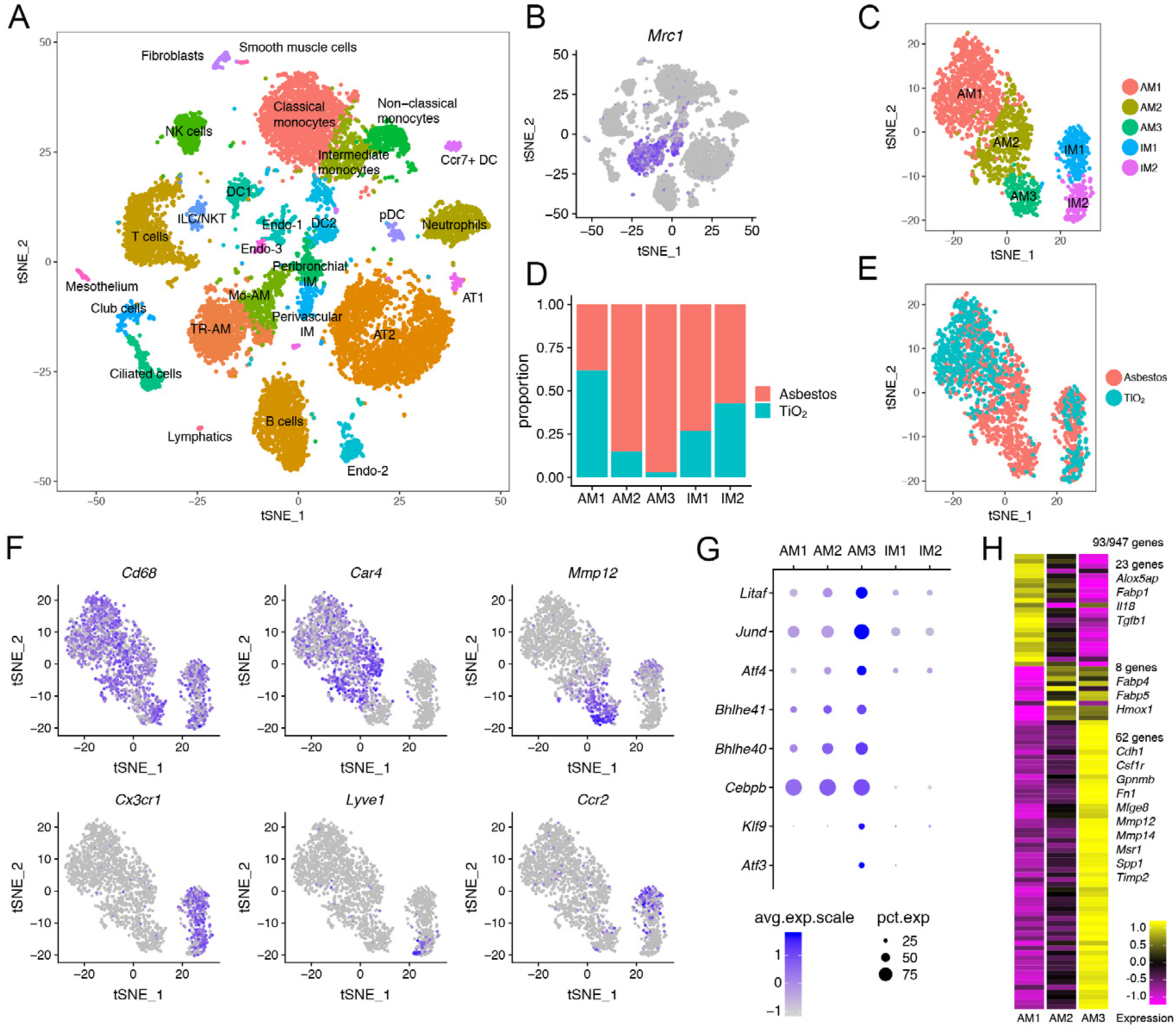
Single-cell RNA-seq reveals profibrotic monocyte-derived alveolar macrophages during asbestos-induced pulmonary fibrosis. **A.** tSNE plot demonstrating 26 cell clusters from 24,060 cells identified by single-cell RNA-seq 14 days after asbestos-or TiO_2_ exposure, one mouse per condition. **B.** Macrophages were identified using canonical lineage-restricted markers, such as *Mrc1*, as shown on a tSNE plot. **C.** Clusters of cells expressing *Mrc1* were subset from the main dataset and re-clustered, revealing two subclusters of tissue-resident interstitial lung macrophages (IM1 and IM2) and three subclusters of alveolar macrophages (AM1, AM2, AM3). **D-E.** Bar plot (**D**) and feature plot (**E**) demonstrating the composition of macrophage subclusters in cells from asbestos and TiO_2_-exposed animals. **F**. Feature plots demonstrating expression of cluster-specific genes: *Cd68* as a pan-macrophage marker, *Car4* as a marker of mature tissue-resident alveolar macrophages (AM1, AM2) and *Mmp12* as a marker of monocyte-derived alveolar macrophages (AM3). Tissue-resident interstitial macrophages are characterized by expression of *Cx3cr1* and can be further subdivided into perivascular (*Lyve1*) and peribronchial (*Ccr2*) interstitial macrophages. **G**. Dot plot demonstrating the expression of transcription factors differentially expressed in monocyte-derived alveolar macrophages (AM3). **H**. Heatmap of 93 genes overlapping between alveolar macrophages and pulmonary fibrosis-associated genes from the Comparative Toxicogenomic Database (947 genes as of February 2019). Selected genes characterizing clusters are shown, see Figure S4E for the full list of genes.

**Figure S4.**
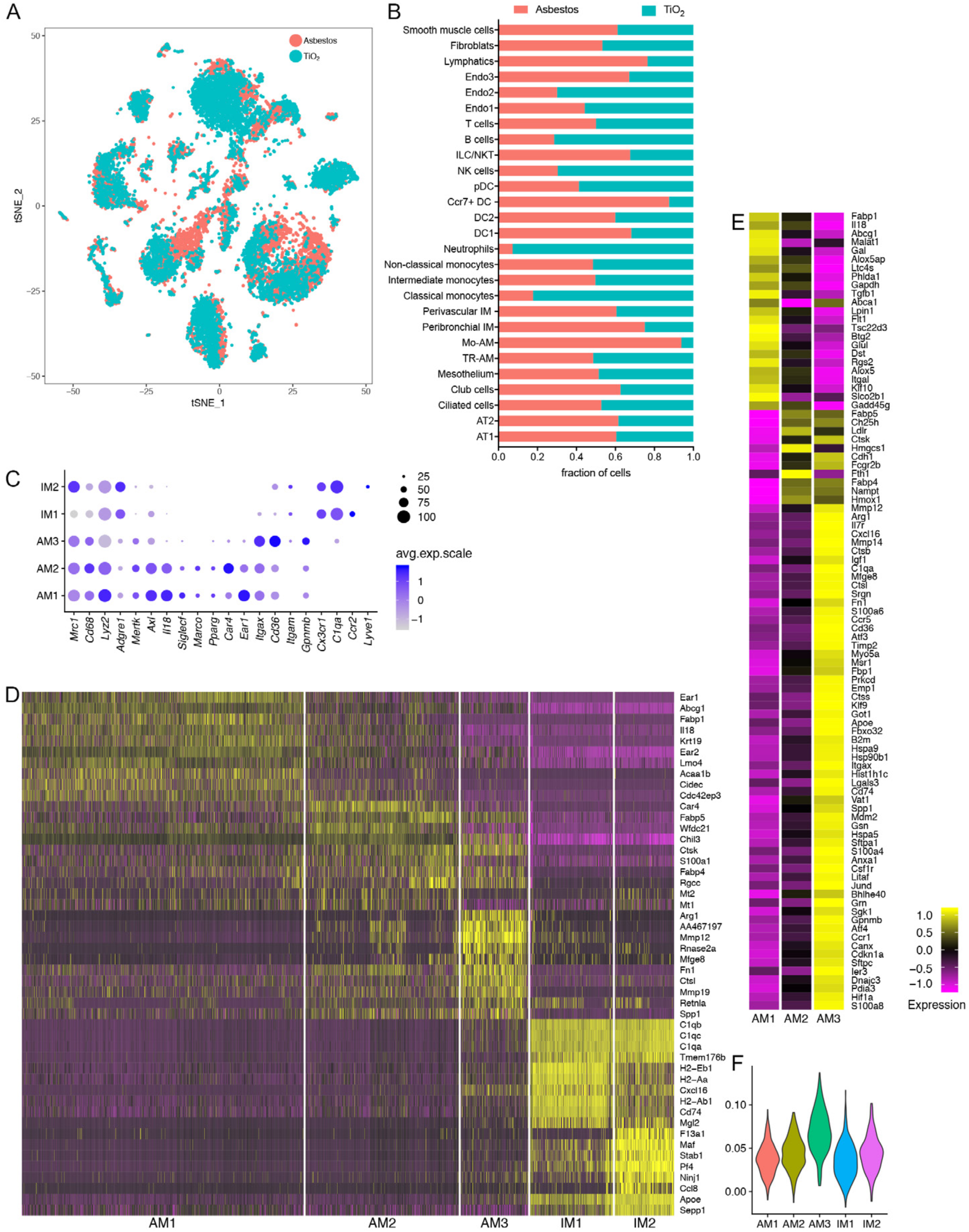
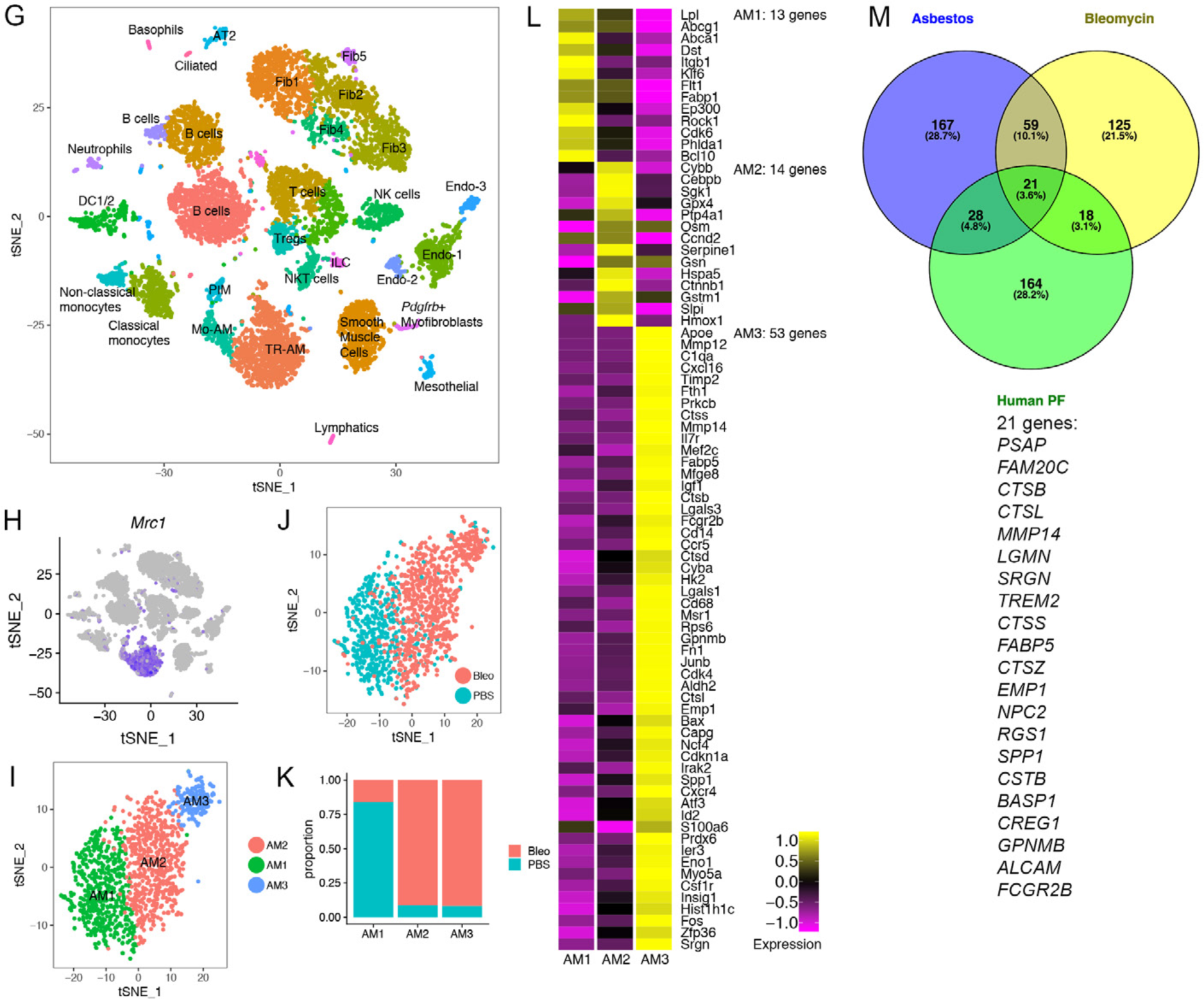
Single-cell RNA-seq identifies heterogeneous response of resident lung cellular populations during asbestos-induced pulmonary fibrosis. (refers to Figure 4). **A-B**. tSNE (**A**) and bar plot (**B**) demonstrating composition of cell clusters. One mouse per library/condition. **C.** Dot plot demonstrating expression of selected genes in macrophage subclusters. **D**. Heatmap of top 10 genes (by log fold change) characterizing macrophage subclusters. **E**. Heatmap of 93 genes overlapping between genes expressed in alveolar macrophages and pulmonary fibrosis-associated genes from the Comparative Toxicogenomic Database (947 genes as of February 2019) (p = 4.561944e-06 for cluster AM3, hypergeometric probability test). **F**. Violin plot demonstrating enrichment of cluster AM3 for the expression of genes associated with pulmonary fibrosis in the Comparative Toxicogenomic Database. The enrichment score was generated using AddModuleScore function as implemented in the Seurat 2.3.4 R toolkit. **G**. tSNE plot demonstrating cell clusters (10,372 cells) identified by single-cell RNA-seq 21 days after PBS or bleomycin treatment, three mice per condition. Data from Xie et al., 2018. **H**. Macrophages were identified using canonical lineage-restricted markers, such as *Mrc1*, as shown on a tSNE plot. **I**. Focused analysis of alveolar macrophages identifies 3 subclusters: AM1, AM2, AM3. **J-K.** tSNE (**J**) and bar plot (**K**) demonstrating composition of alveolar macrophages clusters. **L**. Heatmap showing 80 genes overlapping between genes expressed in alveolar macrophages in Xie et al. dataset and pulmonary fibrosis-associated genes from the Comparative Toxicogenomic Database (947 genes as of February 2019) (p = 3.388409e-05 for cluster AM3, hypergeometric probability test). **M**. Venn diagram showing overlap between the genes in cluster AM3 from asbestos dataset, cluster AM3 in Xie et al. 2018 dataset and genes from the profibrotic macrophage cluster (cluster 1) from Reyfman et al., 2018. List of 21 genes overlapping between three datasets is shown. See also supplementary table S5.

We identified macrophages based on the expression of canonical macrophage-associated genes (*Cd68, Mrc1, Lyz2, Adgre1* and *Axl*) (Figure 4B, Figure S4C). Focused analysis of the macrophage populations resolved two major populations identified as alveolar macrophages and tissue-resident interstitial macrophages. Tissue-resident interstitial macrophages could be further subdivided into peribronchial and perivascular macrophages, distinguished by expression of *Ccr2* and *Lyve1*, respectively (Figure 4C-F, Figure S4C) (Gibbings et al., 2017). While some markers were expressed in both alveolar and interstitial macrophages (*Cd68, Mrc1, Lyz2, Adgre1* and *Axl*), others were restricted to alveolar macrophages (*Siglec F, Marco* and *Il18*) (Figure S4C; Supplemental Table S2).

Alveolar macrophages contained three subclusters. Cluster AM1 was comprised of alveolar macrophages from mice exposed to either asbestos or TiO_2_ (Figure 4D, E). In contrast, clusters AM2 and AM3 were predominantly represented by cells from asbestos-exposed animals. Alveolar macrophages from cluster AM1 were characterized by expression of genes associated with normal homeostatic function of alveolar macrophages (*Ear1, Fabp1*) (Figure S4D). Macrophages from cluster AM2 expressed genes involved in inflammatory response, cytokine production and matrix metalloproteinase activation (*Car4, Ctsk, Chil3, S100a1, Wfdc21*) (Figure S4D). Macrophages from cluster AM3 exhibited a more immature alveolar macrophage phenotype, characterized by lower expression of *Pparg, Car4, Ear1, Siglecf, Marco*, and increased expression of *Itgam, Cd36, Gpnmb* (Figure S4C-D) and increased expression of transcription factors involved in macrophage development and maintenance (*Litaf, Jund, Bhlhe40, Bhlhe41, Klf9*) and unfolded protein response (*Atf3, Atf4*,) (Figure 4G). We queried the expression of 947 genes associated with pulmonary fibrosis using Comparative Toxicogenomic Database (Davis et al., 2017) and found that 93 genes were detected in alveolar macrophages (Figure 4H, Figure S4E). Clusters AM1 and AM2 contained “generic” genes (23 and 8 genes correspondingly) associated with alveolar macrophage cellular identity and function (*Fabp1, Fabp4, Fabp5, Tgfb1*) (Gautier et al., 2012). In contrast, cluster AM3 was enriched for profibrotic genes and contained 62 genes involved in matrix remodeling and cell to cell interactions (*Cdh1, Gpnmb, Fn1, Mmp12, Mmp14, Spp1, Timp2*) (Figure 4H, Figure S4E, F).

We then investigated whether our findings in the model of asbestos-induced pulmonary fibrosis share similarities with a commonly used model of bleomycin-induced pulmonary fibrosis. In a recent publication Xie et al. evaluated the response of fibroblasts in bleomycin-induced pulmonary fibrosis using single-cell RNA-seq (Xie et al., 2018). While the authors focused their analysis on fibroblasts, their dataset contained a large number of non-mesenchymal cells, including alveolar macrophages (Figure S4G-H; Supplemental Table S3). Analysis of the macrophage population demonstrated presence of the three subclusters: AM1, AM2, AM3 (Figure S4I; Supplemental Table S4). Similar to our asbestos dataset, cluster AM1 was predominantly comprised by cells from PBS-treated control animals, while clusters AM2 and AM3 were comprised by cells from bleomycin-treated animals (Figure S4J,K). Out of 947 genes associated with pulmonary fibrosis in the Comparative Toxicogenomic Database we found 80 to be detected in alveolar macrophages from bleomycin dataset (Figure S4L). Cluster AM3 was enriched for expression of fibrosis-associated genes (53 genes), many of which overlapped with the AM3 cluster genes from our asbestos dataset (*Gpnmb, Fn1, Mmp12, Mmp14, Spp1, Timp2*) (Figure S4F-M).

We have previously reported the emergence of a novel subpopulation of alveolar macrophages in lung explants from patients with pulmonary fibrosis compared with biopsies from the normal donor lung (Reyfman et al., 2018). Twenty one genes, including *SPP1, MMP14, TREM2, GPNMB* overlapped between the genes characterizing the cluster of human profibrotic alveolar macrophages from patients with pulmonary fibrosis and genes observed in the AM3 clusters from asbestos-induced and bleomycin-induced pulmonary fibrosis (Figure S4M; Supplementary Table S5). Thus, our analysis has demonstrated the emergence of homologous populations in different models of pulmonary fibrosis (asbestos and bleomycin) and in patients with pulmonary fibrosis.

### Autocrine M-CSF/M-CSFR signaling is necessary for the maintenance of monocyte-derived alveolar macrophages within fibrotic niches

Tissue-resident alveolar macrophages rely on GM-CSF produced by alveolar epithelial cells for their homeostatic maintenance. In contrast, data from M-CSF-deficient mice and pharmacological blockade of M-CSFR suggest that M-CSF signaling is dispensable for homeostatic maintenance of alveolar macrophages (Sauter et al., 2014; Shibata et al., 2001; Witmer-Pack et al., 1993). We found that *Csf1r*, encoding M-CSFR, was upregulated in AM3 cluster both in asbestos-and bleomycin-treated mice (Figure S4E, L). We asked which cells produce M-CSF and IL-34 – ligands for M-CSFR – and particularly whether any new M-CSF/IL-34-producing cells emerge during fibrosis. *Il34* was expressed in alveolar type II cells and club cells in control as well as asbestos-treated animals, while *Csf1* was expressed in fibroblasts, mesothelium and endothelium (Figure S5A,B). Surprisingly, we found *Csf1* expression in AM3 cluster in both asbestos and bleomycin models (Figure 5A, B). Expression of *Csf1* was also detected in bulk RNA-seq from flow-sorted macrophages in our previously published data from the bleomycin model (Figure 5C) (Misharin et al., 2017). We validated our findings using *in situ* RNA hybridization. Double positive *Mrc1*^*+*^*Csf1*^*+*^ alveolar macrophages were detected in the areas of fibrosis in proximity to *Pdgfra*^*+*^ fibroblasts 28 days after asbestos exposure (Figure 5D, E; Figure S5C). Thus, our data suggest that monocyte-derived alveolar macrophages are capable of generating an autocrine signal for their persistence. Consistent with this hypothesis, administration of anti-CSF1 or anti-CSF1R antibodies 14 days after asbestos exposure reduced the number of alveolar macrophages (Figure 5F,G). While expression of *CSF1* was detected in alveolar macrophages from both donor and fibrotic human lung (Figure S5D), expression of *CSF1R* was increased in patients with pulmonary fibrosis, specifically in the cluster containing profibrotic alveolar macrophages (Figure S5E,F).

**Figure 5.**
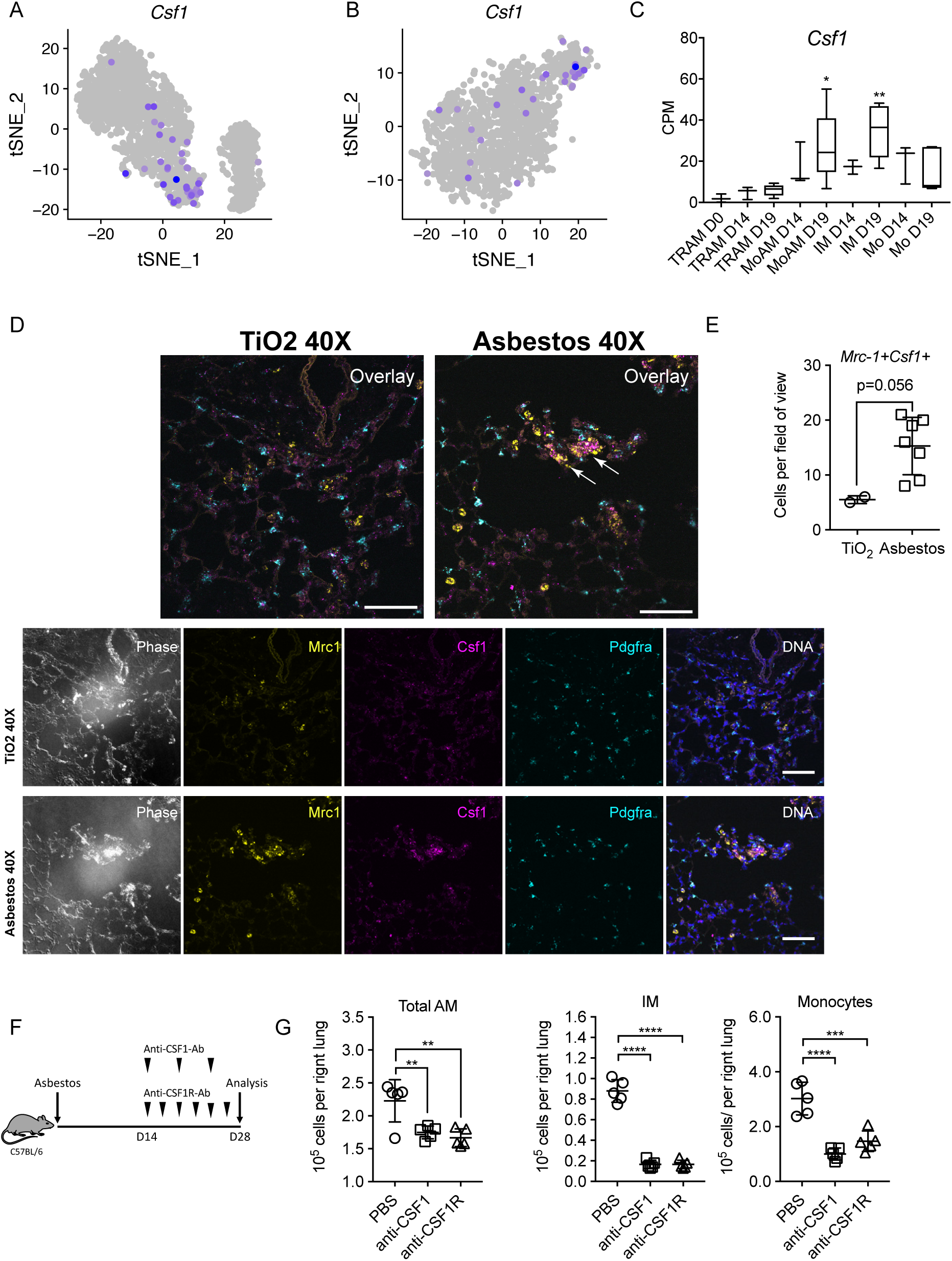
Autocrine M-CSF signaling is required for maintenance of monocyte-derived alveolar macrophages within fibrotic niches. **A-B.** tSNE plot showing expression of *Csf1* in alveolar macrophages after asbestos (**A**) and bleomycin (**B**) exposure. **C**. Barplot showing expression of *Csf1* in flow-sorted alveolar macrophages during the course of bleomycin-induced pulmonary fibrosis. Data from Misharin et al., 2017. One-way ANOVA with Bonferroni correction for multiple comparisons; *, P < 0.05; **, P < 0.01. **D**. *In situ* RNA hybridization confirms expression of *Csf1* in alveolar macrophages during asbestos-induced pulmonary fibrosis. Analysis performed on day 28 post-TiO_2_ or asbestos exposure. Macrophages were identified as *Mrc1*^*+*^, fibroblasts as *Pdgfra*^*+*^ cells. Arrows indicate *Mrc1*^*+*^*Csf1*^*+*^ alveolar macrophages. Scale bar is 50 μm. **E**. Number of *Mrc1*^*+*^*Csf1*^*+*^ alveolar macrophages is increased after asbestos exposure. Data are from 2 mice, mean±SD, Mann-Whitney test. **F.** Schematic of experimental design. C57BL/6 mice were administered crocidolite asbestos (100 μg, intratracheally), treated with anti-CSF1 antibody (intraperitoneally, every 5 days), anti-CSF1R antibody (0.5 mg, intraperitoneally, 3 times a week) or PBS from day 14 to day 28. Number of monocytes and macrophages was measured by flow cytometry at day 28. **G.** Total alveolar macrophages, interstitial macrophages and monocytes from asbestos-exposed animals according to the experimental design in **F**. Data presented as mean±SEM, 5 mice per group, one-way ANOVA with Tukey-Kramer test for multiple comparisons; *, P < 0.05; **, P < 0.01; ***, P < 0.001; ****, P < 0.0001.

**Figure S5.**
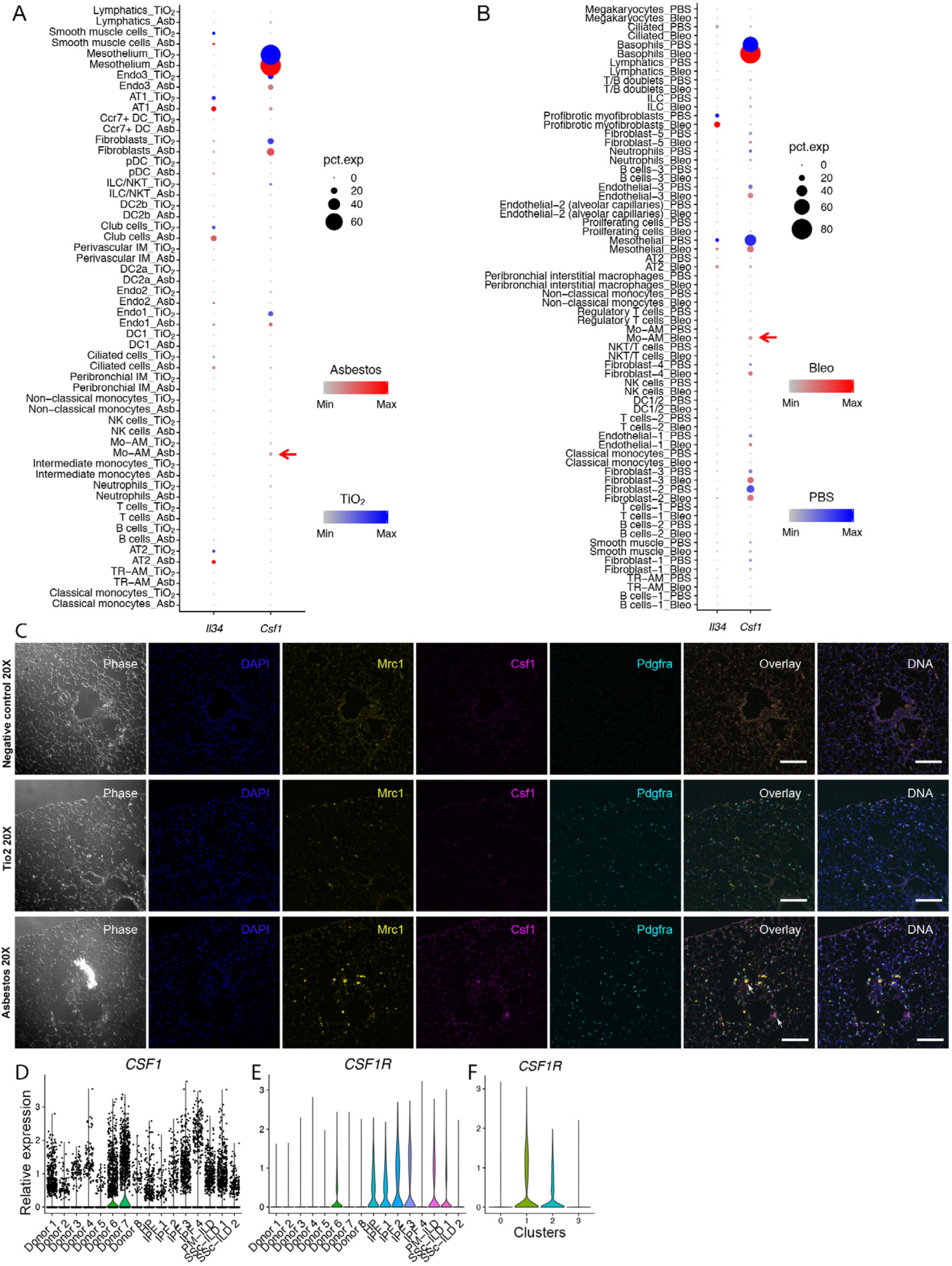
Autocrine M-CSF signaling is required for maintenance of monocyte-derived alveolar macrophages in fibrotic niches (refers to Figure 5). **A-B.** Dot plot showing expression of *Csf1* and *Il34* across cell types during asbestos-(**A**) and bleomycin-induced pulmonary fibrosis (**B**). Arrows point out expression of *Csf1* in monocyte-derived alveolar macrophages. **C**. *In situ* RNA hybridization confirms expression of *Csf1* in alveolar macrophages during pulmonary fibrosis. Analysis performed on day 28 post-TiO_2_ or asbestos exposure. Top row: negative controls (no primary probe), demonstrating level of autofluorescence in the tissue. Scale bar is 50 μm. **D**. *CSF1* is expressed in human alveolar macrophages in all 8 donor lungs and 8 lungs from patients with pulmonary fibrosis. **E-F**. Expression of *CSF1R* is increased in alveolar macrophages from patients with pulmonary fibrosis, specifically, in cluster 1, containing profibrotic alveolar macrophages. Data from Reyfman et al. 2018.

### Monocyte-derived alveolar macrophages provide a link between epithelial injury and activation of resident fibroblasts within spatially-restricted profibrotic niches

We queried both our asbestos and published bleomycin (Xie et al., 2018) alveolar macrophage single-cell RNA-seq datasets for the expression of ligands that have partnering receptors on fibroblasts and could be involved in their proliferation. Using curated database of the ligand-receptor pairs (Ramilowski et al., 2015) we identified that macrophages from AM3 cluster in both asbestos-and bleomycin-treated mice expressed *Pdgfa* (Figure 6A, B). We confirmed increased expression of *Pdgfa* in monocyte-derived alveolar macrophages by querying our bulk RNA-seq data from flow-sorted macrophages from the bleomycin model (Figure 6C) (Misharin et al., 2017). Finally, we validated our findings by performing *in situ* RNA hybridization for *Mrc1, Pdgfa, Pdgfra* after asbestos exposure and detected double positive *Mrc1*^*+*^*Pdgfa* ^*+*^ alveolar macrophages in the areas of fibrosis in proximity to *Pdgfra*^*+*^ fibroblasts (Figure 6D, E; Figure S6).

**Figure 6.**
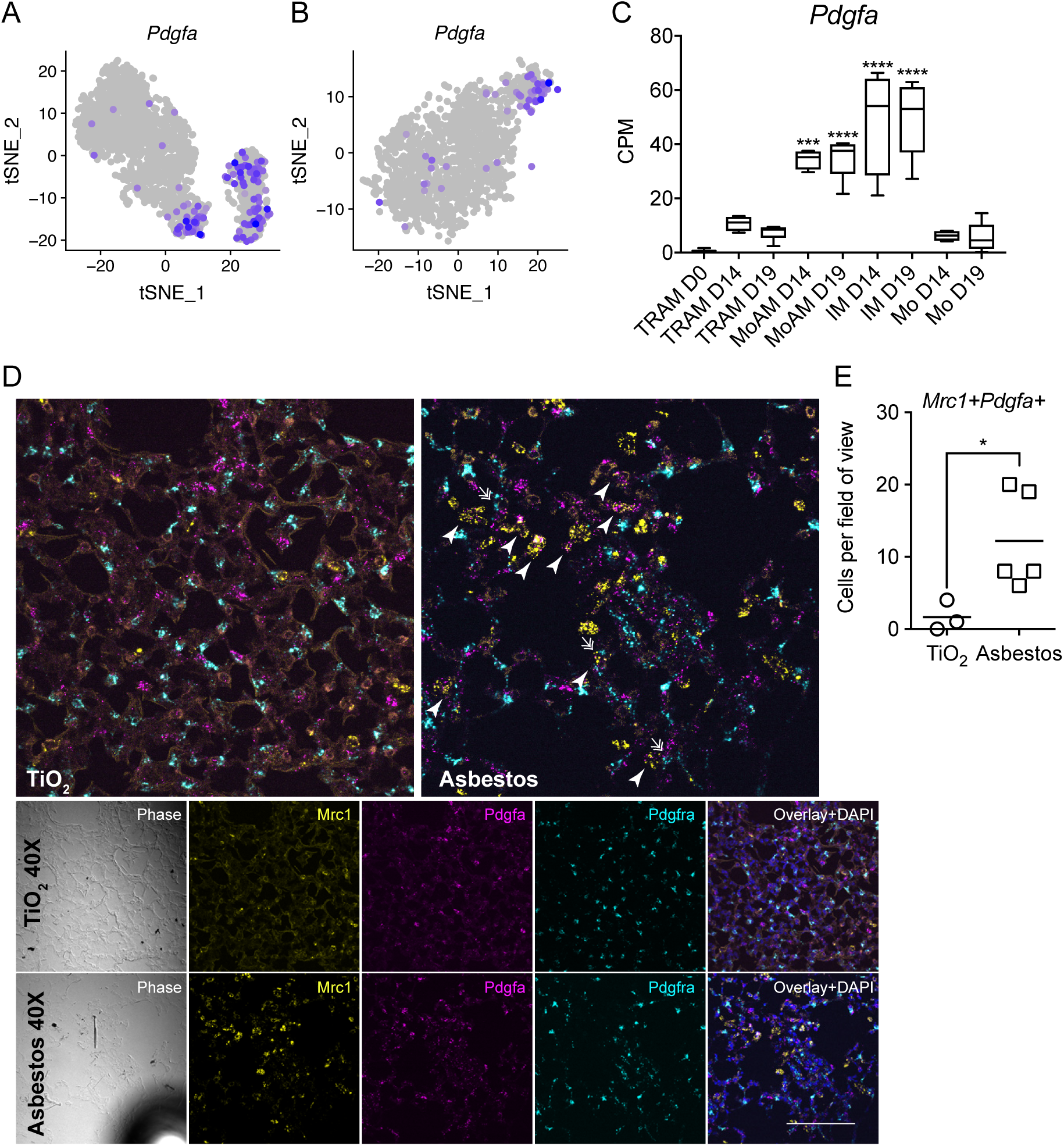
Monocyte-derived alveolar macrophages express *Pdgfa* which is required for fibroblast proliferation. **A-B.** tSNE plot showing expression of *Pdgfa* in alveolar macrophages after asbestos (**A**) and bleomycin (**B**) exposure. **C**. Barplot showing expression of *Pdgfa* in flow-sorted alveolar macrophages during the course of bleomycin-induced pulmonary fibrosis. Data from Misharin et al., 2017. One-way ANOVA with Bonferroni correction for multiple comparisons; ***, P < 0.001, ****, P < 0.0001. **D**. *In situ* RNA hybridization confirms expression of *Pdgfa* in alveolar macrophages during pulmonary fibrosis. Analysis performed on day 28 post-TiO_2_ or asbestos exposure. Macrophages were identified as *Mrc1*^*+*^, fibroblasts as *Pdgfra*^*+*^ cells. Arrow-heads indicate *Mrc1*^*+*^*Pdfga*^*+*^ macrophages, double arrows indicate *Pdgfra*^*+*^ fibroblasts adjacent to *Mrc1*^*+*^*Pdfga*^*+*^ macrophages. Scale bar is 50 μm. **E**. Number of *Mrc1*^*+*^*Pdgfa*^*+*^ alveolar macrophages is increased after asbestos exposure *, P < 0.05. Data are from 2 mice, mean±SD, Mann-Whitney test.

**Figure S6.**
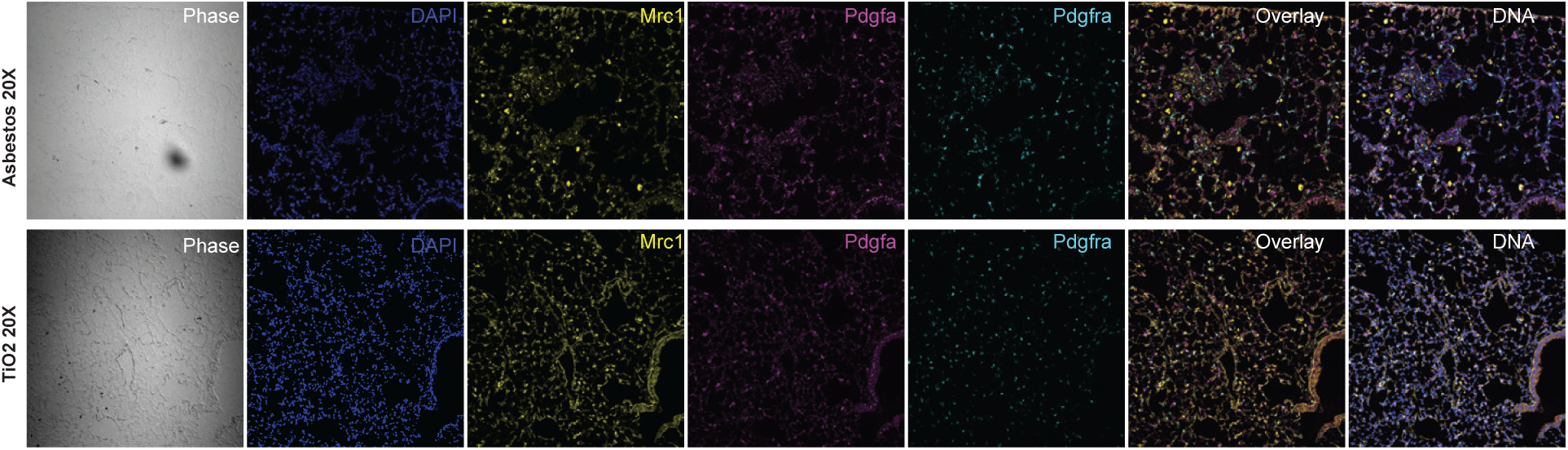
Monocyte-derived alveolar macrophages express *Pdgfa* which is required for fibroblast proliferation. (refers to Figure 6). *In situ* RNA hybridization confirms expression of *Pdgfa* in alveolar macrophages during pulmonary fibrosis, 20x magnification. Analysis performed on day 28 post-TiO_2_ or asbestos exposure. Scale bar is 50 μm.

To explore the possibility that monocyte-derived alveolar macrophages represent a link between asbestos-induced epithelial injury and fibroblast activation we performed analysis of alveolar epithelial type II cells, which were well-represented in our dataset. We did not detect distinct subclusters within alveolar type II cells. However, *Retnla* (RELMα) was differentially expressed in alveolar type II cells between asbestos and TiO_2_-exposed animals (Figure 7A; Supplemental Table S6). *Retnla* is primarily expressed in epithelial cells and is thought to act as a chemoattractant for monocytes and monocyte-derived cells. Moreover, *Retnla*-deficient mice have been reported to be protected from experimental pulmonary fibrosis (Liu et al., 2014). We hypothesized, that in a model of spatially-restricted pulmonary fibrosis, expression of *Retnla* will be restricted to areas of fibrosis. We treated *Cx3cr1*^*ER-Cre*^*ZsGreen* mice with tamoxifen 14 and 15 days after administration of asbestos or TiO_2_ and analyzed lungs using confocal immunofluorescent microscopy at day 21. Double-staining for RELMα and surfactant protein C confirmed that alveolar type II cells expressing RELMα were spatially restricted to areas of fibrosis, which also contained GFP^+^ monocyte-derived alveolar macrophages (Figure 7B). Thus, our data suggest that monocyte-derived alveolar macrophages link spatially-restricted epithelial injury and fibroblast proliferation.

**Figure 7.**
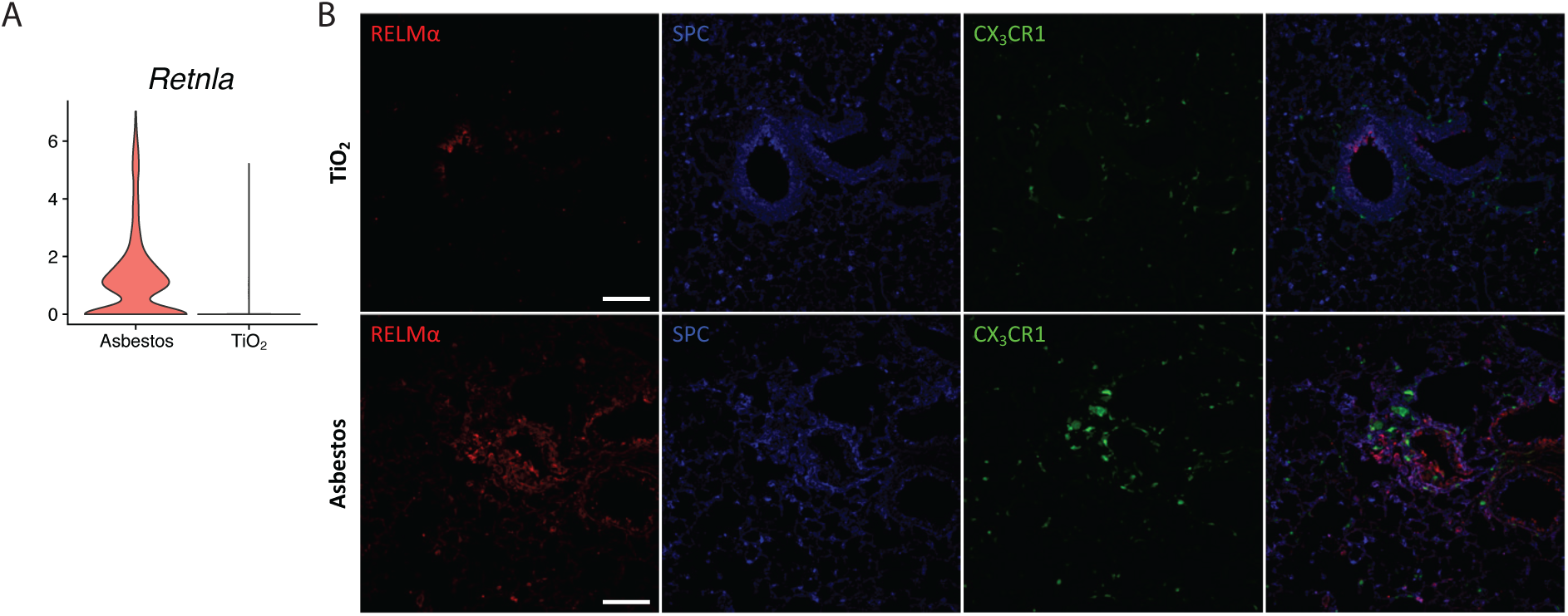
Expression of RELMα is restricted to epithelial cells located in the areas of fibrosis. **A.** Violin plot demonstrating increased expression of *Retnla* in alveolar type II cells 14 days after asbestos exposure. Log fold change 2.83, p value 1.00E-166, Wilcoxon rank-sum test. **B.** Representative fluorescent images showing expression of RELMα (red), SPC (blue), and CX_3_CR1-GFP (green) in lungs from TiO_2_-or asbestos-treated animals at day 21 post-exposure. RELMα is detected in the airway epithelial cells and alveolar type II cells in the fibrotic regions in asbestos model, but not in alveolar type II cells after TiO_2_ exposure. Scale bar 100 μm.

## Discussion

We demonstrate that while tissue-resident interstitial macrophages, tissue-resident alveolar macrophages, and monocyte-derived alveolar macrophages are present in the fibrotic niche, only monocyte-derived alveolar macrophages are causally related to fibrosis. Moreover, we demonstrate that single-cell RNA-seq combined with spatial methods, such as *in situ* RNA hybridization and immunofluorescence microscopy, can be used to identify mechanisms of intracellular communication within the fibrotic niche. Specifically, we predicted that monocyte-derived alveolar macrophages are capable of self-maintenance via autocrine M-CSF/M-CSFR signaling, and confirmed this *in vivo*. Furthermore, increased expression of *Retnla* on epithelial cells, *Pdgfa* on monocyte-derived alveolar macrophages and *Pdgfra* in fibroblasts emerged from analysis of single-cell RNA-seq data and could be localized to the fibrotic niche, thereby generating testable hypotheses about the mechanisms of the intercellular interactions necessary for fibrosis.

Single-cell RNA-seq is transforming our understanding of biology by revealing heterogeneity within cell populations that emerge during disease. We show that single-cell RNA-seq can be used to combine data generated by different laboratories in different animal models of disease with homologous data from diseased patients to identify common mechanisms in disease pathogenesis (Reyfman et al., 2018; Xie et al., 2018). By using a genetic lineage tracing system that marks the ontogeny of recruited macrophages, we were able to spatially localize intercellular signals predicted from single-cell RNA-seq data to areas of fibrosis surrounding asbestos fibers in the mouse lung. Computational consolidation of single-cell RNA-seq datasets generated in other models of fibrosis is a promising approach to generate hypotheses with respect to fibrosis pathogenesis that can be tested with causal genetic or pharmacologic interventions (Lotfollahi et al., 2018).

Our findings suggest the emergence of spatially restricted multicellular pro-fibrotic niches during pulmonary fibrosis. In these niches, injured epithelial cells drive recruitment of monocyte-derived alveolar macrophages, which, in turn, provide signals for fibroblast proliferation. Tissue-resident macrophages rely on growth factors produced by other cells comprising the niche (Guilliams and Scott, 2017), particularly on M-CSF produced by fibroblasts (Zhou et al., 2018), or in case of tissue-resident alveolar macrophages, on GM-CSF, which is normally produced by alveolar type II cells (Guilliams et al., 2013). These self-regulating circuits between different cell types are essential for the maintenance of tissue-resident macrophages during homeostasis, as was recently demonstrated in an elegant study of a two-cell system containing macrophages and fibroblasts (Zhou et al., 2018). In contrast, we found that monocyte-derived alveolar macrophages and profibrotic alveolar macrophages in patients with pulmonary fibrosis were characterized by increased expression of both *Csf1* and *Csf1r* suggesting they can maintain their population in the fibrotic niche via autocrine production of M-CSF independent of signals from other resident lung cells. This may provide a mechanism by which macrophages are sustained at the “advancing front” of fibroblastic foci where myofibroblasts are actively proliferating and matrix remodeling is occurring (Herrera et al., 2019). In this niche profibrotic macrophages secrete factors essential for fibroblast proliferation, including PDGFA, which has been shown to be elevated in patients with pulmonary fibrosis (Martinet et al., 1987; Nagaoka et al., 1990). This might also offer an opportunity for therapy targeting M-CSF/M-CSFR signaling as was recently shown to be effective in the model of the radiation-induced fibrosis (Meziani et al., 2018). Furthermore, while monocyte-derived alveolar macrophages recruited to the lung during bleomycin also express M-CSF and M-CSFR, bleomycin-induced fibrosis resolves spontaneously. Combining our lineage tracing system with single-cell RNA-seq data from resolving bleomycin-induced fibrosis and persistent asbestos-induced fibrosis might suggest how monocyte-derived alveolar macrophages change as fibrosis resolves.

Using previously validated genes as markers, we were able to use our single-cell RNA-seq data to resolve three transcriptionally and anatomically distinct populations of macrophages in the normal murine lung—alveolar macrophages, peribronchial interstitial macrophages and perivascular interstitial macrophages, which is in agreement with previously published studies (Gibbings et al., 2017; Lim et al., 2018). These data further suggested emergence of a fourth macrophage population, monocyte-derived alveolar macrophages, during fibrosis, which we were able to confirm with a genetic lineage tracing system. Moreover, we identified the emergence of a novel cell state within tissue-resident alveolar macrophages after asbestos exposure. These data highlight the importance of genetic lineage tracing studies and single-cell spatial transcriptomics or proteomics to validate putative cell populations identified from RNA-seq data.

The differentiation of circulating monocytes into alveolar macrophages is a complex process that involves reshaping of nearly 70% of the cellular transcriptome (Lavin et al., 2014). This requires the coordinated action of multiple signaling pathways—the disruption of any of which could prevent the differentiation of monocytes into alveolar macrophages. In this context, protection against fibrosis in macrophage-specific knockouts of a given gene might result from a failure of monocyte to alveolar macrophage differentiation. For example, monocyte/alveolar macrophage deletion of *Tgfb1* (Yu et al., 2017), *Torc1* (Sinclair et al., 2017), *Pparg* (Schneider et al., 2014), *Casp8* (Misharin et al., 2017) or *Cflar* (McCubbrey et al., 2018) prevents monocyte to macrophage differentiation, likely explaining their salutary effects on fibrosis or airway inflammation.

Our study has limitations. First, while we used a validated lineage tracing system to exclude the possibility that tissue-resident peribronchial or perivascular interstitial macrophages serve as a significant source of profibrotic alveolar macrophages, it is possible that they play an important role in responding to tissue injury. Second, while our single-cell RNA-seq analysis was able to capture and resolve many of the canonical cell types identified in the mouse lung, several cell types, including alveolar type I cells, fibroblasts and smooth muscle cells/myofibroblasts were underrepresented in our dataset. Thus, we were not able to resolve heterogeneity within these populations during pulmonary fibrosis. Other techniques, such as enrichment for specific populations of interest or single-nucleus RNA-seq, may address these limitations (Wu et al., 2019; Xie et al., 2018; Zepp et al., 2017).

In summary, we found that the recruitment of monocyte-derived alveolar macrophages is limited to spatially restricted fibrotic niches in pulmonary fibrosis. Genes expressed by injured epithelial cells, monocyte-derived macrophages and fibroblasts can be detected using single-cell RNA-seq and spatially organized using spatial transcriptomics. Genetically inducing the deletion of monocyte-derived alveolar macrophages by necroptosis during their differentiation ameliorated lung fibrosis, causally linking them to disease pathogenesis. Single-cell RNA-seq suggested these cells sustain themselves via autocrine signaling through M-CSF/M-CSFR, and the administration of an antibodies to M-CSF or M-CSFR resulted in their loss. Collectively, our results support the combination of lineage tracing, computational synthesis of single-cell RNA-seq datasets from murine and human fibrosis and *in situ* RNA hybridization imaging as a powerful method to identify pathways that can be targeted for the treatment of different disease endotypes in patients with pulmonary fibrosis.

## Supporting information

Supplemental_Table_S1_all_clusters

Supplemental_Table_S2_Asbestos_Macrophages

Supplemental_Table_S3_Xie_Bleo_all_cells

Supplemental_Table_S4_Xie_Bleo_Macrophages

Supplemental_Table_S5_profibrotic_macrophages_overlap

Supplemental_Table_S6_AT2_TiO2vsAsbestos

## Materials and Methods

### Mice

All mouse procedures were approved by the Institutional Animal Care and Use Committee at Northwestern University (Chicago, IL, USA). All strains including wild-type mice are bred and housed at a barrier-and specific pathogen–free facility at the Center for Comparative Medicine at Northwestern University. Ten to twelve week old mice were used for all experiments. The C57BL/6J, *Cx3cr1*^*ER-Cre*^ mice (Yona et al., 2013) and ZsGreen (Madisen et al., 2010) mice were obtained from Jackson laboratories (Jax stocks 000664, 020940 and 007906, correspondingly). *Casp8*^*flox/flox*^, *CD11c*^*Cre*^*Casp8*^*flox/flox*^ and *CD11c*^*Cre*^*Casp8*^*flox/flox*^*RIPK3*^*-/-*^ mice have been described previously (Misharin et al., 2017).

### Asbestos-induced lung fibrosis and drug administration

Eight to ten week old mice were instilled intratracheally with control particles (TiO_2_ [100 µg in 50 µl PBS]) or crocidolite asbestos fibers (100 µg in 50 µl PBS) to induce lung-fibrosis, as previously described (Cheresh et al., 2015). For lineage-tracing studies, anesthetized mice were administered tamoxifen by oral gavage [100 µl of 10 mg tamoxifen (Sigma, T5648, St. Lois, Missouri) dissolved in sterile corn oil (Sigma, C8267, St. Lois, Missouri]. Anti-CSF1 antibody (clone BE0204, BioXCell; 0.5mg, intraperitoneally, every 5 days) or anti-CSF1R antibody (clone BE0213, BioXCell; 0.5mg, intraperitoneally, 3 times a week) are used *in vivo*, as previously described (Naik et al., 2015; Ruffell et al., 2014). Lungs were harvested at indicated time points for flow cytometry, single-cell RNA sequencing, histopathology, immunohistochemistry and immunofluorescence.

### Tissue preparation and flow cytometry

Tissue preparation for flow cytometry analysis and cell sorting was performed as previously described (Misharin et al., 2017), with modifications. Briefly, mice were euthanized and their lungs were perfused through the right ventricle with 10 ml of HBSS. The lungs were removed and infiltrated with 2 mg/ml collagenase D (Roche, Indianapolis, Indiana) and 0.2 mg/ml DNase I (Roche, Indianapolis, Indiana) dissolved in HBSS with Ca^2+^ and Mg^2+^, using syringe with 30G needle. Lungs were chopped with scissors, tissue was transferred into C-tubes (Miltenyi Biotech, Auburn, California, 130-096-334), and processed in a GentleMACS dissociator (Miltenyi Biotech, Auburn, California) using m_lung_01 program, followed by incubation for 30 minutes at 37°C with gentle agitation, followed by m_lung_02 program. The resulting single-cell suspension was filtered through a 40 μm nylon cell strainer mesh to obtain a single-cell suspension. The cells were incubated with anti-mouse CD45 microbeads (Miltenyi Biotech, Auburn, California,130-052-301) and CD45^+^ cells were collected using the MultiMACS™ Cell24 Separator (Miltenyi Biotech, Auburn, California), according to the manufacturer’s protocol. Automated cell counting was performed (Nexcelom K2 Cellometer with AO/PI reagent). Cells were stained with fixable viability dye eFluor 506 (eBioscience, Waltham, Massachusetts), incubated with FcBlock (BD Biosciences, San Jose, California), and stained with a following mixture of fluorochrome-conjugated antibodies (listed as antigen, clone, fluorochrome, manufacturer, catalog number): MHC II, 2G9, BUV395, BD Biosciences, 743876; Ly6C, HK1.4, eFluor450, eBiosciences, 48-5932-82; CD45, 30-F11, FITC, eBiosciences, 11-0451-82; CD64, X54-5/7.1, PE, Biolegend, 139303; Siglec F, E50-2440, PECF594, BD Biosciences, 562757; CD11c, HL3, PECy7, BD Biosciences, 558079; CD24, M1/69, APC, eBiosciences, 17-0242-82; CD11b, M1/70, ACPCy7, Biolegend, 101225; Ly6G, 1A8, Alexa 700, BD Biosciences, 561236; NK1.1, PK136, Alexa 700, BD Biosciences, 560515. Single color controls were prepared using BD CompBeads (BD Biosciences, San Jose, California) and Arc beads (Invitrogen, Waltham, Massachusetts). Flow cytometry was performed at the Northwestern University Robert H. Lurie Comprehensive Cancer Center Flow Cytometry Core facility (Chicago, Illinois). Data were acquired on a custom BD FACSymphony instrument using BD FACSDiva software (BD Biosciences, San Jose, California). Compensation and analysis were performed using FlowJo software (TreeStar). Each cell population was identified using sequential gating strategy (Figure S1A). The percentage of cells in the live/singlets gate was multiplied by the number of live cells using Cellometer K2 Image cytometer to obtain an cell count.

### Single-cell RNA-seq

Single-cell suspensions were prepared as described above with slight modification. Mice were euthanized with sodium pentobarbital overdose. Chest cavity was opened and lungs were perfused through the right ventricle with 10 ml of HBSS. The lungs were removed and, using 30G needle, infused with 1 mL dispase (Corning) with DNase I (Sigma). Lungs were incubated at room temperature with gentle agitation for 45 minutes, followed by gentle teasing using forceps into small (1-2 mm) fragments and incubation in digestion buffer for another 15 minutes. Resulting suspension was passed through passed through 70 µm cell strainer (Falcon), washed with DMEM (Corning) supplemented with 5% FBS (Corning), pelleted by centrifugation and erythrocytes were lysed using BD Pharm Lyse (BD Biosciences). Resulting single cell suspension was kept in DMEM/FBS and passed through 40 µm cell strainer (Falcon) two times. Cells were counted using Cellometer K2 (Nexcelom) with nucleic acid binding dyes acridine orange (AO) to calculate total number of nucleated cells and propidium iodide (PI) to count dead cells, cell viability exceeded 85%. All manipulations were performed using wide bore tips (Axygen). Single-cell 3’ RNA-Seq libraries were prepared using Chromium Single Cell V2 Reagent Kit and Controller (10x Genomics, Pleasanton, California). Libraries were assessed for quality (TapeStation 4200, Agilent, Santa Clara, California) and then sequenced on HiSeq 4000 instrument (Illumina, San Diego, California), for single-cell RNA-Seq libraries. Initial data processing was performed using the Cell Ranger version 2.0 pipeline (10x Genomics, Pleasanton, California). Analysis was performed using the Seurat R toolkit V2.3.4 (Butler et al., 2018) and R 3.4, see supplemental materials for detailed R code.

### Data availability

Single-cell RNA-seq data from TiO_2_-and asbestos-exposed mice have been deposited to GEO (GSE127803). We also used bulk RNA-seq data on flow-sorted alveolar macrophages from GSE82158 (Misharin et al., 2017), single-cell RNA-seq data from patients with pulmonary fibrosis GSE122960 (Reyfman et al., 2018) and mice exposed to bleomycin GSE104154 (Xie et al., 2018).

### Fluorescence *in situ* RNA hybridization

Multiplex fluorescent *in situ* hybridization was performed using RNAscope (Advanced Cell Diagnostics (ACD), Newark, CA). Mouse lungs were inflated to 15 cm of H_2_O and fixed with 4% paraformaldehyde (EMS) for 24 h. Lungs were paraffin embedded, and 5 μm tissue sections were mounted on Superfrost Plus slides (Fisher, Waltham, Massachusetts). Slides were baked for 1 h at 60°C, deparaffinized in xylene and dehydrated in 100% ethanol. Sections were treated with hydrogen peroxide (ACD 322381) for 10 min at room temperature, then heated to mild boil (98-102°C) in 1x target retrieval reagent buffer (ACD 322001) for 15 min. Protease Plus (ACD 322381) was applied to sections for 30 min at 40°C in HybEZ Oven (ACD). Hybridization with target probes, preamplifier, amplifier, fluorescent labels and wash buffer (ACD 320058) was done according to ACD instructions for Multiplex Fluorescent Reagent Kit v2 (ACD 323100) and 4-Plex Ancillary Kit v2 (ACD 323120) when needed. Parallel sections were incubated with ACD positive (ACD 321811) and negative (ACD 321831) control probes. Sections were covered with ProLong Gold Antifade mountant (Thermo, Waltham, Massachusetts, P36930). Probes used were: mouse *Mrc1* (ACD 437511-C3), mouse *Pdgfa* (ACD 411361), mouse *Pdgfra* (ACD 480661-C2) and mouse *Csf1* (ACD 315621). Opal fluorophores (Opal 520 (FP1487001KT), Opal 620 (FP1495001KT), Opal 690 (FP1497001KT), Perkin Elmer, Shelton, CT) were used at 1:1500 dilution in Multiplex TSA buffer (ACD 322809). Images were captured on Nikon A1R confocal microscope with 20x and 40x objectives. Images were uniformly processed and pseudocolored in Fiji.

### Histopathology, immunohistochemistry and immunoflourescence

For histopathology and immunohistochemistry, mouse lung tissue from the regions adjacent to the regions used for flow cytometry and single-cell RNA-Seq was fixed in 4% paraformaldehyde for 24 hours, dehydrated and embedded in paraffin. 4 µm thick sections were prepared. Hematoxylin and eosin staining and Masson’s trichrome staining were performed for the analysis of fibrosis scoring. Immunohistochemistry was performed at Northwestern University Mouse Histology and Phenotyping Laboratory Core facility (Chicago, Illinois).

For immunofluorescence, mouse lung tissue was fixed in 4% paraformaldehyde for 6 hours and transferred into 20% sucrose for overnight incubation. Tissue was embedded in Tissue-Tek OCT compound (Sakura, Torrance, California), flash frozen in liquid nitrogen and cut on cryostat at 14 µm thickness. Sections were air-dried and stained with PE-conjugated anti-MERTK (BioLegend, San Diego, California, 151505), Alexa Fluor 647-conjugated anti-Siglec F (BD Biosciences, San Jose, California, 562680), rabbit anti-RELMα (Abcam, Cambridge, Massachusetts, ab39626), and goat anti-SPC (Santa Cruz Biotechnology, Dallas, Texas, sc-7706). Appropriate secondary antibodies were used for unconjugated primary antibody, including donkey anti-rabbit Alexa Fluor 647 (Invitrogen, Waltham, Massachusetts, A-31573) and donkey anti-goat Alexa Flour 568 (Invitrogen, Waltham, Massachusetts, A-11057). DAPI (Invitrogen, D3571) was used for nuclear staining and sections were mounted with ProLong Diamond Antifade Mountant (Invitrogen, Waltham, Massachusetts, P36965). Images were acquired on Nikon A1R confocal microscope or Nikon Ti2 wide field microscope at Northwestern University Nikon Cell Imaging Facility (Chicago, Illinois), processed using Nikon Elements Software.

### Fibrosis Scores and Lung Collagen Determination

Fibrosis and fibrosis scores in mice were determined from hematoxylin and eosin and Masson’s trichrome–stained specimens in a blinded manner, in accordance with the code set by Pathology Standards for Asbestosis, as described previously (Kamp et al., 1995). Collagen levels were determined using Sircol assay as described previously (Cheresh et al., 2015).

## Statistical analysis

Statistical tests and tools for each analysis are explicitly described with the results or detailed in figure legends.

## Acknowledgments

This work was supported by the Office of the Assistant Secretary of Defense for Health Affairs, through the Peer Reviewed Medical Research Program under Award W81XWH-15-1-0215 to Drs. Budinger and Misharin. Opinions, interpretations, conclusions and recommendations are those of the author and are not necessarily endorsed by the Department of Defense. Next generation sequencing on the Illumina HiSeq 4000 was performed by the NUSeq Core Facility, which is supported by the Northwestern University Center for Genetic Medicine, Feinberg School of Medicine, and Shared and Core Facilities of the University’s Office for Research. Northwestern University Flow Cytometry Facility, Center For Advanced Microscopy, and Pathology Core Facility are supported by NCI Cancer Center Support Grant P30 CA060553 awarded to the Robert H.Lurie Comprehensive Cancer Center. Multiphoton microscopy was performed on a Nikon A1R multiphoton microscope, acquired through the support of NIH 1S10OD010398-01. This research was supported in part through the computational resources and staff contributions provided by the Genomics Computing Cluster (Genomic Nodes on Quest) which is jointly supported by the Feinberg School of Medicine, the Center for Genetic Medicine, and Feinberg’s Department of Biochemistry and Molecular Genetics, the Office of the Provost, the Office for Research, and Northwestern Information Technology.

## Author contributions

Nikita Joshi: designed the study, performed experiments, analyzed results, performed bioinformatics analysis, wrote manuscript. Satoshi Watanabe: designed the study, performed experiments, analyzed results, wrote manuscript. Rohan Verma: performed bioinformatics analysis, wrote manuscript. Renea P. Jablonski, Ching-I Chen, Paul Cheresh: performed experiments, analyzed results. Paul A. Reyfman: performed bioinformatics analysis. Alexandra C. McQuattie-Pimentel, Lango Sichizya, Annette S. Flozak: performed experiments, analyzed results. Cara J. Gottardi: analyzed results, wrote manuscript. Carla M. Cuda, Harris Perlman: developed and provided genetically modified animals, provided reagents and resources. Manu Jain: provided reagents and resources, wrote manuscript. David Kamp: designed and supervised the study, provided reagents and resources, wrote manuscript. GR Scott Budinger and Alexander V.Misharin: designed and supervised the study, performed analysis, wrote manuscript, provided funding for the project. All authors discussed the results and commented on the manuscript.

